# Substrate-induced assembly and functional mechanism of the bacterial membrane protein insertase SecYEG-YidC

**DOI:** 10.1101/2025.05.26.656142

**Authors:** Max Busch, Cristian Rosales-Hernandez, Michael Kamel, Yulia Schaumkessel, Eli O. van der Sluis, Otto Berninghausen, Thomas Becker, Roland Beckmann, Alexej Kedrov

## Abstract

The Sec translocon and the YidC/Oxa1-type insertases universally mediate biogenesis of α-helical membrane proteins, but the molecular basis of their cooperation has remained disputed. Recent discoveries of multi-subunit insertases assembled at the back of the translocon in fungi and higher eukaryotes have raised the question about the architecture and mechanism of the putative bacterial ortholog SecYEG-YidC. Here, we combine cryogenic electron microscopy with cell-free protein synthesis to visualize biogenesis of SecYEG/YidC-dependent multipass membrane protein NuoK. We demonstrate that the nascent chain of NuoK does not enter the lateral gate of SecYEG, but crosses over the translocon towards its back side, whereto YidC is recruited in the nascent chain-dependent manner. The SecY-YidC interface promotes folding of the transmembrane helices prior their insertion, in agreement with thermodynamic principles of membrane protein folding. YidC forms extensive contacts with the nascent chain, suggesting its key role in the insertion event. Our data provide detailed insights on the insertase machinery, suggest the evolutionary conservation of the gate-independent insertion route, and offer an expanded view on membrane protein biogenesis.

## Introduction

Integral membrane proteins (IMPs) are encoded by ∼30 % protein-coding genes in each organism and they determine the functionality of the cellular membranes (von Heijne, 2007). Biogenesis of α-helical IMPs at the cytoplasmic membrane of bacteria and the endoplasmic reticulum (ER) in eukaryotes typically occurs in a co-translational manner: The nascent transmembrane helices (TMHs) emerging from the ribosome are identified by the signal recognition particle (SRP) and delivered in the form of ribosome-nascent chain complexes (RNCs) to the universally conserved Sec translocon that facilitates their insertion in the lipid bilayer (Cymer *et al*, 2015b; Itskanov & Park, 2023). The translocon of *Escherichia coli* is a heterotrimeric complex composed of SecY, SecE and SecG proteins forming a central pore. N-and C-terminal halves of the central subunit SecY may open laterally in a clam-shell manner, resulting in a lipid-exposed gap between the opposing TMHs 2b and 7, located at the front side of the translocon. This gap, commonly referred as the “lateral gate”, has been seen as the exclusive site for the nascent TMHs to partition into the lipid phase, both for bacterial SecYEG and the homologous Sec61αβγ in eukaryotes. This model has been supported by extensive biochemical studies (du Plessis *et al*, 2009; Sachelaru *et al*, 2013) and via structural insights, commonly focused on signal peptides of secretory proteins and single-pass IMPs exposing their N-terminal ends into the cytoplasm (Jomaa *et al*, 2016; Kater *et al*, 2019; Voorhees & Hegde, 2016). However, multipass IMPs have their TMHs in alternating topology, suggesting higher complexity within the biogenesis pathways (Bischoff *et al*, 2014; Ou *et al*, 2025).

In bacteria, biogenesis of multiple IMPs relies on a cooperative action of SecYEG and the essential and ubiquitous membrane protein insertase YidC (Steinberg *et al*, 2018; Zhu *et al*, 2013). To facilitate folding and membrane insertion of the nascent IMPs, YidC forms a polar water-filled groove in the cytoplasmic lipid leaflet, and its unusually short TMHs 3, 4 and 5 may distort the lipid bilayer (Chen *et al*, 2017; McDowell *et al*, 2021). The groove between TMHs 3 and 5 in the core of the membrane serves for insertion of the client IMPs (Kedrov *et al*, 2016; Wickles *et al*, 2014). Extensive intermolecular crosslinking studies suggested that YidC is located in front of the lateral gate of SecYEG where it could access the nascent TMHs emerging from the lateral gate (Jauss *et al*, 2019; Petriman *et al*, 2018; Sachelaru *et al*., 2013), and the model was further supported by low-resolution cryo-electron microscopy (cryo-EM) analysis (Botte *et al*, 2016). Remarkably, the AlphaFold algorithm confidently predicts a very different architecture for the machinery of *E. coli*, with YidC placed at the back of SecYEG, and the docking is barely affected by presence of other SecYEG-associated proteins, such as SecDF and PpiD-YfgM (Fig. EV1 and Appendix Fig. S1) (Miyazaki *et al*, 2022; Tsukazaki, 2018). Nearly identical SecYEG-YidC assemblies are predicted for other γ-proteobacteria, including pathogens *Pseudomonas aeruginosa* and *Klebsiella pneumoniae*, while diverse architectures are proposed for species from other bacterial clades (Appendix Fig. S1D; Appendix Table S1, Dataset EV2). Notably, the confidence of the models beyond γ-proteobacteria is low, suggesting that other factors, such as accessory proteins, specific lipids, but also the bound ribosome and the nascent chain may be required for the complex assembly.

Recent biochemical and high-resolution cryo-EM studies of the eukaryotic insertase machinery revealed that the YidC homolog TMCO1 was located at the back of the Sec61 complex, being a component of the large multi-subunit complex BOS-GEL-PAT, also referred as multipass membrane protein translocon (MPT) (Smalinskaite *et al*, 2022; Sundaram *et al*, 2022). Furthermore, insertion of a nascent IMP opsin occurred not via the lateral gate, but between Sec61 and the BOS complex, challenging the long-standing paradigm in the field. Complementary, global ribosome profiling has suggested that the MPT is assembled in a substrate-dependent manner, and that hundreds of eukaryotic multipass IMPs utilize the “back-of-Sec” insertion route (Sundaram *et al*, 2025). The accumulated controversy between the insights from the eukaryotic homologs and the conventional model of the SecYEG-YidC complex has raised the question about the mechanism of IMP folding in bacteria and the architecture and dynamics of the insertion machinery (Smalinskaite & Hegde, 2023). To tackle this issue, here we employed nanodisc-reconstituted SecYEG-YidC and cell-free protein synthesis (CFPS) to analyze the biogenesis of the bacterial multipass IMP NuoK, the subunit K of the NADH-quinone oxidoreductase that requires both components for the membrane integration and folding in *E. coli* (Price & Driessen, 2010). By means of cryo-EM, we describe the detailed architecture of the RNC-bound SecYEG-YidC complex at different statges of NuoK insertion. YidC is recruited to the back of SecYEG in the substrate-dependent manner thus indeed resembling the settings at the eukaryotic insertase machinery. The nascent chain emerging from the ribosome folds within a hydrophobic pocket formed by SecY loops and then it is routed away from the lateral gate to be inserted between SecYEG and YidC. Overall, the data elucidate the functional architecture of the bacterial cotranslational IMP insertion machinery and reveal the evolutionary conservation of the “back-of-Sec” insertion route, thereby offering a new perspective on membrane protein biogenesis in bacteria.

## Results

### YidC engages the nascent chain of NuoK at a late stage of insertion

Though multiple membrane proteins require both SecYEG and YidC for their insertion and folding, only for few of them the biogenesis pathways have been investigated in detail. In particular, mass spectrometry of YidC-depleted *E. coli* membranes indicated that NuoK, a subunit of the respiratory complex I (Fig. 1A and Appendix Fig. S2A), is dependent on the insertase (Price & Driessen, 2008). *In vitro* experiments showed that YidC was required for insertion of NuoK TMH 2 and TMH 3 in proteoliposomes (Fig. 1B) (Price & Driessen, 2010). We questioned whether interactions of YidC with the nascent NuoK occured in the cellular membrane. We assembled defined co-translational insertion intermediates of NuoK *in vivo*, where N-terminal fragments of NuoK containing either 48 or 86 residues (NuoK^48^ and NuoK^86^) were followed by a 30 aa long linker, including haemeagglutinine (HA)-tag and a C-terminal ribosome stalling sequence derived from SecM (here referred to as SecM*; Fig. 1C and Appendix Fig. S1B) (Cymer *et al*, 2015a; Kempf *et al*, 2017). With this construct, the SecM* peptide and the linker/tag would occupy the exit tunnel of the stalled ribosome, while two or three NuoK TMHs would be exposed for NuoK^48^ and NuoK^86^, respectively, being accessible for the insertase complex. After expressing the constructs and lysing the cells, the isolated membranes were solubilized with detergent and subjected to centrifugation through sucrose cushion, so only large macromolecular complexes, such as ribosomes and the associated proteins, were expected to be sedimented. Immunodetection confirmed equal amounts of YidC in all membrane preparations, while presence of YidC in the cushion pellets was clearly specific to the nascent chain (Fig. 1D and Appendix Fig. S3D): Only faint signals were detected in absence of NuoK or upon expression of NuoK^48^, while NuoK^86^ strongly promoted recruitment of YidC. Thus, we concluded that YidC engages the nascent NuoK at rather late insertion stage *in vivo*, which can be recapitulated in a reconstituted system (Price & Driessen, 2010).

**Figure 1.**
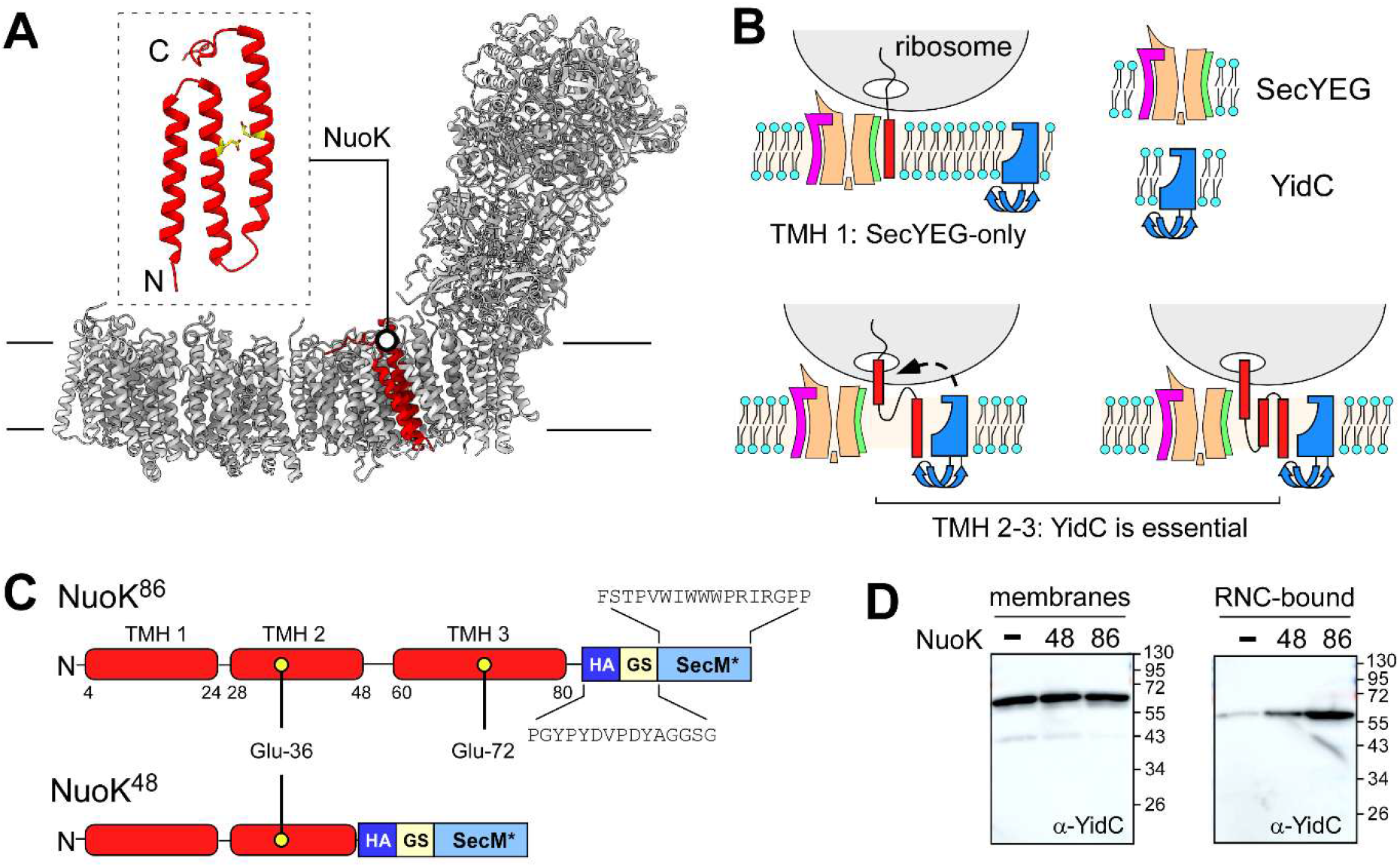
YidC insertase is involved in *E. coli* NuoK biogenesis. **(A)** Structure of the *E. coli* NuoK subunit within the assembled respiratory complex I (PDB ID 7NYR). Glu-36 and Glu-72 residues are shown in yellow. **(B)** Presumable insertion pathway of NuoK. Insertion of NuoK TMH 1 is mediated by SecYEG, while YidC is required for NuoK TMH 2 and TMH 3. **(C)** Design of the nascent chains mimicking NuoK^48^ and NuoK^86^ insertion intermediates. NuoK TMHs are shown in red with the flanking residue positions indicated. Positions of the glutamate residues within TMHs 2 and 3 are shown as yellow circles. The sequence of the C-terminal linker consisting of the HA-tag (HA) and a glycine-serine linker (GS) is shown for NuoK^86^. SecM* represents a SecM-based ribosome stalling sequence. **(D)** Immunodetection of YidC in *E. coli* membranes (left) and the ribosome-bound material (right) upon expression of the NuoK insertion intermediates. The NuoK length is indicated above.

### SecYEG and YidC-dependent insertion in nanodiscs

While our efforts to visualize the SecYEG-YidC complex in detergent by means of cryo-EM were not successful, possibly due to destabilized protein:protein interactions in absence of lipids, we set out to elucidate its architecture using lipid-based nanodiscs, a well-defined system for studying IMP insertion (Kater *et al*., 2019; Kedrov *et al*, 2013; Kedrov *et al*., 2016; Koch *et al*, 2018) (Fig. 2A). To co-reconstitute both SecYEG and YidC in the relevant topology and to avoid their stochastic distribution among the nanodiscs, YidC and SecE were genetically fused via a cleavable linker (Appendix Fig. S3A), while the non-essential cationic C-terminal end of YidC was removed to exclude spontaneous electrostatic interactions with the ribosome observed *in vitro* (Kedrov *et al*., 2013; Kohler *et al*, 2009). The fusion protein was expressed and isolated together with SecY and SecG subunits (Fig. 2B and Appendix Fig. S3B). The complex was reconstituted into a sufficiently large nanodisc formed by the scaffold protein MSP2N2 (expected diameter up to 17 nm) (Ritchie *et al*, 2009) and treated with the HRV-3C protease to release YidC from SecE (Fig. 2C and Appendix Fig. 3C-E). As a result, each nanodisc contained one copy of both SecYEG and YidC, presumably in the correct orientation, while the dimensions of the surrounding lipid bilayer would allow for lateral diffusion of the proteins, assembly of the functional complex and insertion of the nascent TMHs. Supporting this notion, the molecular weight distribution of the nanodisc-embedded SecYEG-YidC complexes measured by single-molecule mass photometry peaked at 290 kDa (Fig. 2D), and the difference from the calculated total protein weight of ∼225 kDa likely reflected the presence of the co-reconstituted lipids. The lipid membrane consisted of 70 mol % POPC and 30 mol % POPG, thus mimicking the ratio of zwitterionic/anionic lipids and the degree of the acyl chain unsaturation found within the bacterial membrane (Sohlenkamp & Geiger, 2016).

**Figure 2.**
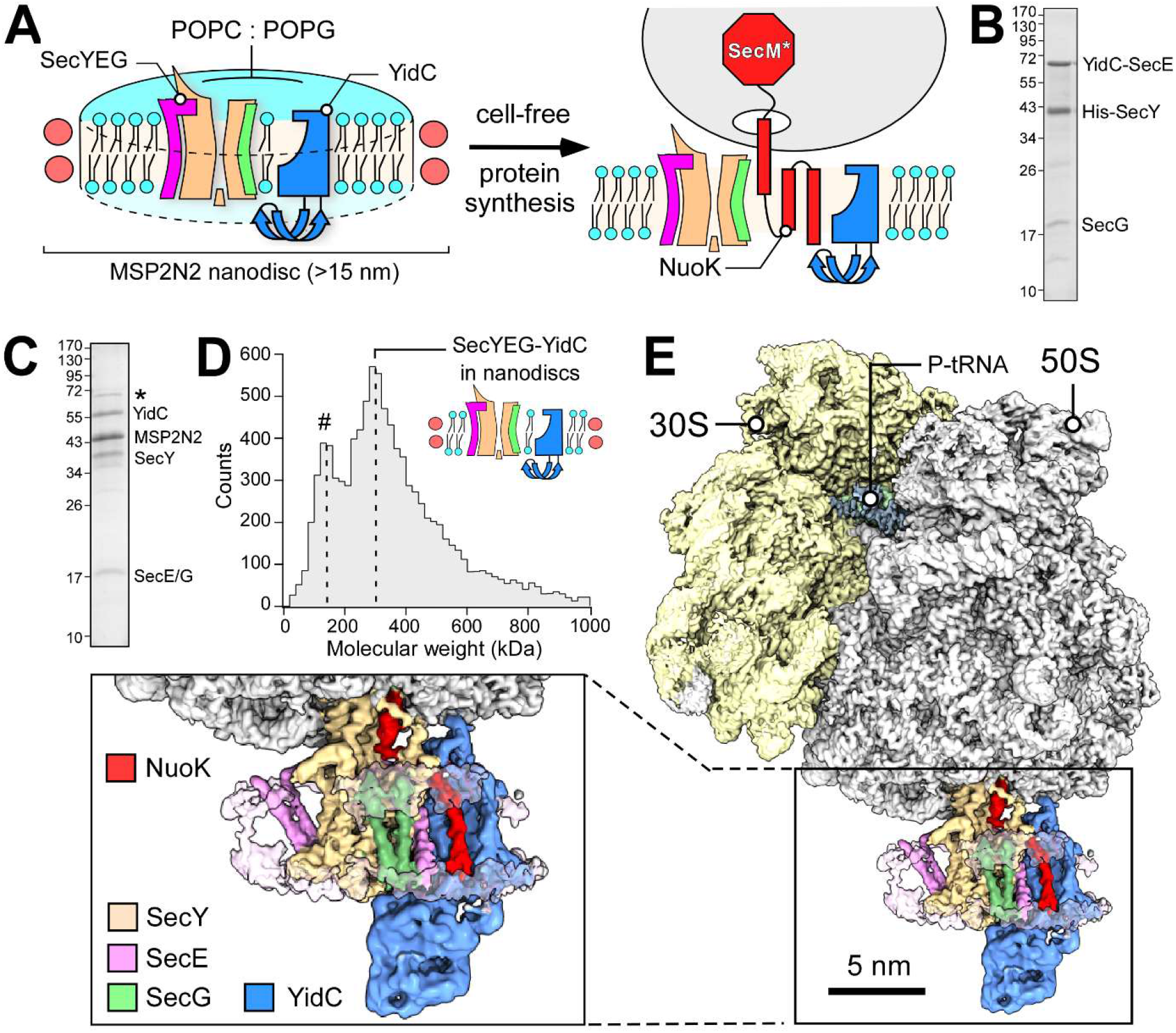
Substrate-induced assembly of the RNC:SecYEG-YidC complex. **(A)** Scheme of the assay to study NuoK biogenesis. SecYEG and YidC are co-reconstituted into MSP2N2-based nanodisc in presence of POPC.POPG lipids and introduced into CFPS reaction. Synthesis of the substrate NuoK is interrupted by SecM* stalling sequence, so the stable insertion intermediate is formed for structural analysis. **(B)** SDS-PAGE of the affinity-purified SecYEG-YidC fusion insertase complex, with the subunits indicated. **(C)** SDS-PAGE of the nanodisc-reconstituted SecYEG-YidC complex after the protease treatment. The minor band at 70 kDa (*) corresponds to the residual YidC-SecE fusion protein. **(D)** Mass photometry recording of the isolated SecYEG-YidC nanodiscs manifests the main peak at 290 kDa. The minor peak at 130 kDa (#) corresponds to the nanodiscs loaded only with lipids. **(E)** Cryo-EM reconstruction of NuoK^86^-RNC:SecYEG-YidC assembly. Shown is a composite map consisting of isolated densities of the ribosome subunits 30S and 50S, P-site peptidyl-tRNA (“P-tRNA”) and locally refined SecYEG-YidC, with subunits and the substrate NuoK indicated.

**Figure 3.**
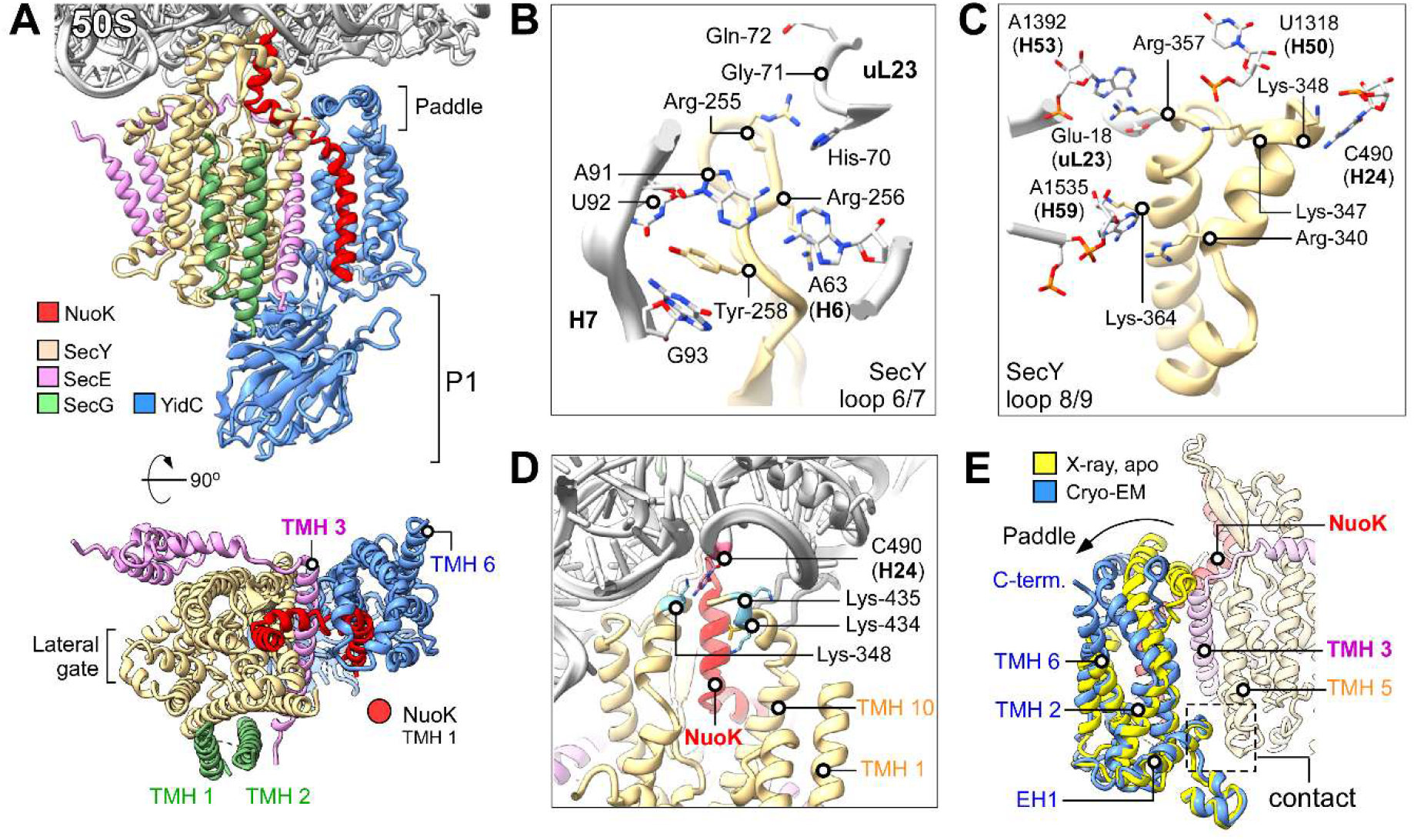
Architecture of the SecYEG-YidC insertase and its interactions with the RNC. (**A**) Molecular model of the SecYEG-YidC insertase complex bound to NuoK^86^-RNC. The putative position of the membrane-inserted NuoK TMH 1 is indicated as a red circle in the cytoplasmic view (bottom). (**B, C**) The network of ribosome:SecYEG interactions mediated by SecY cytoplasmic loops 6/7 and 8/9. The ribosomal RNA and proteins are shown in gray. **(D)** The C-terminal extansion of SecY contacts the flipped-out base C490 of 23S rRNA. This interaction is likely stabilized by Lys-435 close to the H24 rRNA backbone and Lys-348 stacking on the C490 from the opposite site. **(E)** Overlay of X-ray and cryo-EM structures of YidC highlight the conformational change within the insertase. The movement of the YidC paddle domain upon entry of the NuoK nascent chain is indicated with an arrow. The closest SecY-YidC contact at the periplasmic side is indicated with the dashed box. SecY, SecE and NuoK are shown as semi-transparent ribbons for clarity. The crystal structure of *E. coli* YidC resolved in lipidic cubic phase was used as a reference (PDB ID 3WVF).

### Visualization of the active RNC:insertase complex

SecYEG-YidC nanodiscs were supplied to the CFPS reaction based on *E. coli* S30 extract. With that, the translating ribosomes were targeted to SecYEG-YidC, presumably by the endogenous SRP and the SRP receptor FtsY, to initiate nascent IMP insertion. The CFPS approach was chosen to ensure sequential incorporation of nascent NuoK TMHs into the membrane and reduce the risk of erroneous insertase:substrate complexes. We chose NuoK^86^ construct that engages with YidC *in vivo* (Fig. 1D) and requires both SecYEG and YidC for insertion in proteoliposomes (Price & Driessen, 2010). Notably, synthesis of NuoK TMH 3 *in vivo* is followed by completion of the translation, so the membrane insertion of the helix occurs post-translationally. Presence of the SecM* stalling sequence at the C-terminal end of the nascent chain in the reconstituted system should trap NuoK TMH 3 exposed from the ribosome, with TMH 1 and TMH 2 already inserted in the membrane, thus enabling visualization of the late-stage folding intermediate and the insertase machinery by cryo-EM.

The NuoK^86^-RNC:SecYEG-YidC complexes were sequentially purified via centrifugation in the sucrose gradient (to isolate 70S ribosomes) and affinity chromatography against poly-histidine-tagged MSP2N2 (to isolate the ribosome-bound nanodiscs), and presence of the stalled nascent chains was conformed via immunoblotting (Appendix Fig. S2C). The samples were stabilized by rapid glutaraldehyde crosslinking before vitrification and cryo-EM analysis. 3D classification of 70S ribosomes followed by focused classification and local refinement of the exit tunnel region revealed two reconstructions containing extra density for SecYEG and YidC (YidC is recognized by its large periplasmic domain P1) and two reconstructions with only SecYEG (Figure EV2A). From the best resolved SecYEG-YidC class we obtained a final density map with an average resolution of 2.4 Å (Fig. EV2B). The reconstruction showed a ribosome programmed with tRNAs in the A- and P-sites, a continuous nascent chain density ranging from the CCA-end of P-site tRNA through the exit tunnel, and a clear density below the tunnel accounting for SecYEG and YidC embedded into a nanodisc of approx. 15 nm in diameter (Fig. 2E). Local refinement of the nanodisc density resulted in a local resolution of 3 to 5 Å for SecYEG and 4 to 7 Å for most parts of YidC (Fig. EV2C) and allowed unambiguous docking of the modelled SecYEG-YidC complex with only minor adjustments (Appendix Fig. S4 and S5). The structure shows the RNC-bound SecYEG in the center of the nanodisc, while YidC is positioned at the back side of SecYEG near SecE TMH 3, with the large P1 domain of YidC exposed to the “periplasmic” side of the nanodisc. As anticipated, an additional density accounting for TMH 2 and TMH 3 of NuoK is found between SecYEG and YidC, building extensive interactions with both proteins within the membrane and at the interface (Fig. 2E). At lower contour levels, an additional transmembrane density is visualized near NuoK TMH 2, which likely represents NuoK TMH 1 (Appendix Fig. S4F), though YidC TMH 1 could not be excluded. Notably, the membrane-inserted fragment appears in an inverted topology in comparison to the full-length NuoK within the assembled complex I, and the possible scenarios for the topology are provided in the Discussion chapter. Importantly, the visulaized route of the nascent polypeptide via the SecYEG-YidC insertase may be taken by multiple other clients, so we believe that the mechanistic insights described below are of general importance for understanding the membrane protein biogenesis in bacteria.

As YidC was reconstituted into the nanodiscs as a fusion protein with SecE TMH 1, it likely was initially positioned near the lateral gate of SecYEG. This arrangement is indeed predicted by AlphaFold for the fusion protein but with high error values for SecYEG:YidC interactions (Fig. EV3A and B), indicating that YidC positioning at the lateral gate is rather unlikely or unstable. After the linker was cleaved, YidC must have relocated substantially within the lipid-filled nanodisc to interact with the client NuoK at the back of SecYEG. Indeed, the observed dimensions of the nanodisc provide sufficient space for the lateral diffusion (Fig. EV3C). Though the confinement could potentially affect the embedded protein assembly, we observed no contacts of either YidC or SecYEG with the edges of the nanodisc, suggesting that the complex was not perturbed by the environment. To test whether the localization of YidC was not artificially induced by the chemical crosslinking, we carried out cryo-EM analysis of the non-crosslinked NuoK^86^-RNC:SecYEG-YidC sample (Appendix Fig. S6). Though somewhat weaker and less resolved, the YidC density was clearly present in the same position at the back of SecYEG, implying that the crosslinked sample reflects the undisturbed architecture of the substrate-bound SecYEG-YidC machinery.

### Architecture of the SecYEG-YidC insertase

The clear density for all TMHs of SecY, SecE and SecG present in the locally refined map offered the by-now best-resolved view on the translocon structure in the lipid environment (Fig. 3A; Appendix Fig. S4). As observed previously for SecYEG-only assemblies, SecY is anchored via its cytoplasmic loops 6/7 and 8/9 to the exit tunnel of the large ribosomal subunit 50S (Fig. 3B and C). Interestingly, SecY loop 6/7 is somewhat remodeled compared to the previously reported RNC-FtsQ:SecYEG assembly (Kater *et al*., 2019). To avoid a clash with the NuoK nascent chain, the loop shifts away from the exit tunnel towards the RNA helix H6/7 and uL23 (Appendix Fig. S7A). Here, SecY Tyr-258 is tightly accommodated in a pocket formed by bases A91, U92 and G93 of the RNA helix H7, while Arg-256 stacks on A63 of H6, and Arg-255 interacts with the loop of uL23 (His-70/Gly-71/Gln-72) (Fig. 3B). In fact, this distortion in the SecY loop 6/7 leads to a re-positioning of SecYEG with respect to the ribosome, with the 50S subunit tilting by ∼10° towards the N-terminal half of SecYEG (Appendix Fig. S7B). Upon tilting, Lys-348 of the SecY loop 8/9 established a contact with the flipped-out base C490 of 23S rRNA (helix H24), and Arg-357 formed a stacking interaction with A1392 (H53). The latter may be further stabilized by a salt bridge with Glu-18 of the proximate ribosomal protein uL23 (Fig. 3C). Further interactions include the backbone of SecY Lys-347 with U1318 (H50), and Arg-340 with Lys-364 approaching the flipped-out base A1535 of H59. The C-terminal cytoplasmic extension of SecY TMH 10 is bowed at Met-425/Ser-426 toward the SecY loop 8/9, so another contact is formed with the flipped-out base C490 of rRNA H24 (likely via Ala-436) (Fig. 3D). Shortening of the polypeptide and deletion of the proximate Lys-433 and Lys-434 appeared lethal for the cells (Chiba *et al*, 2002), highlighting the importance of this poorly studied element of SecYEG. As this SecY:ribosome contact was absent in earlier structures (Bischoff *et al*., 2014; Jomaa *et al*., 2016; Kater *et al*., 2019), we speculated that its formation is associated with the specific nascent chain routed for insertion. Notably, the lateral gate formed by TMHs 2b and 7 is in a tightly closed conformation (Appendix Fig. S4B and C) indicating that it is not employed by the NuoK nascent chain, and the central pore is sealed by the “plug” domain, TMH 2a of SecY.

The conformation of YidC within the RNC-bound complex clearly deviates from the structure of isolated YidC obtained via X-ray crystallography (Kumazaki *et al*, 2014b) (Fig. 3E and Appendix Fig. S4E). The paddle domain, i.e. the hairpin built of helices CH1 and CH2 at the cytoplasmic interface, shifts by 1 nm in direction of YidC TMH 5, and so to the periphery of the complex. There, it appears in close vicinity to the ribosomal protein uL24, but does not form a direct contact (Fig. 3A). The shift of the paddle domain allows the passage of the nascent chain (Fig. 3E). Furthermore, YidC TMH 6 is located at the periphery of the nanodisc, and its C-terminal extension is not required for the ribosome:SecYEG-YidC assembly, though this polypeptide is essential for binding to an RNC in absence of SecYEG *in vitro* (Kedrov *et al*., 2013), and it is also important for the homologous ribosome:Oxa1 interactions in mitochondria (Haque *et al*, 2010). The large density at the periplasmic side matches the structure of the YidC P1 domain known from crystallography studies (Kumazaki *et al*., 2014b; Ravaud *et al*, 2008). With respect to SecYEG, YidC is oriented with its TMH 3 facing TMH 3 of SecE, and the loop of the paddle domain (residues 397-406) faces the N-terminal end of SecY. While the SecY N-terminal polypeptide (residues 1-15) is not resolved in the structure, it may reach the paddle domain, thus explaining the efficient crosslinks observed between the residue 399 of YidC and SecY in *E. coli* membranes (Petriman *et al*., 2018). At the periplasmic side, the essential amphipathic helix EH1 and SecY TMH 5 (Fig. 3E) form the only pronounced contact between these two proteins (discussed below). Apart from this contact, SecYEG and YidC are held together mainly by the NuoK^86^ nascent chain, i.e., TMH 2 within the membrane region and TMH 3 at the cytoplasmic interface, suggesting that YidC is likely recruited to the RNC-bound SecYEG in a substrate-dependent manner.

### Routing and insertion of the nascent NuoK

The nascent chain could be unambiguously traced from the CCA-end of P-site tRNA to the tunnel exit site within the ribosome (Fig.s 4A and B). All side chains of the SecM* stalling element (_111_WWWPRIRGPP_120_) were resolved with Pro-120 attached to the CCA-end of the A-site tRNA, while the preceding modified motif _117_RGP_119_ (_163_RAG_165_ in SecM) was coupled to the CCA-end of the P-site tRNA (Fig. 4B). The conformation of the stalling peptide, tRNAs and critical bases in the peptidyl-transferase center were highly similar to the structure of SecM-stalled *E. coli* ribosome (Gersteuer *et al*, 2024), indicating the same trapping mechanism as described for the `RAG/P` motif and its modified versions. Further, residues Arg-115 and Arg-117 interacted with ψ2504 and A2062, and the aromatic rings of the tryptophan motif (_112_WWW_114_) with bases A2059 and U2586 in the 23S rRNA (Fig. 4C). Less resolved density accounting for the linker and the HA-tag continues from the constriction by uL4 and uL22 towards the mouth of the tunnel (Fig. 4A).

**Figure 4.**
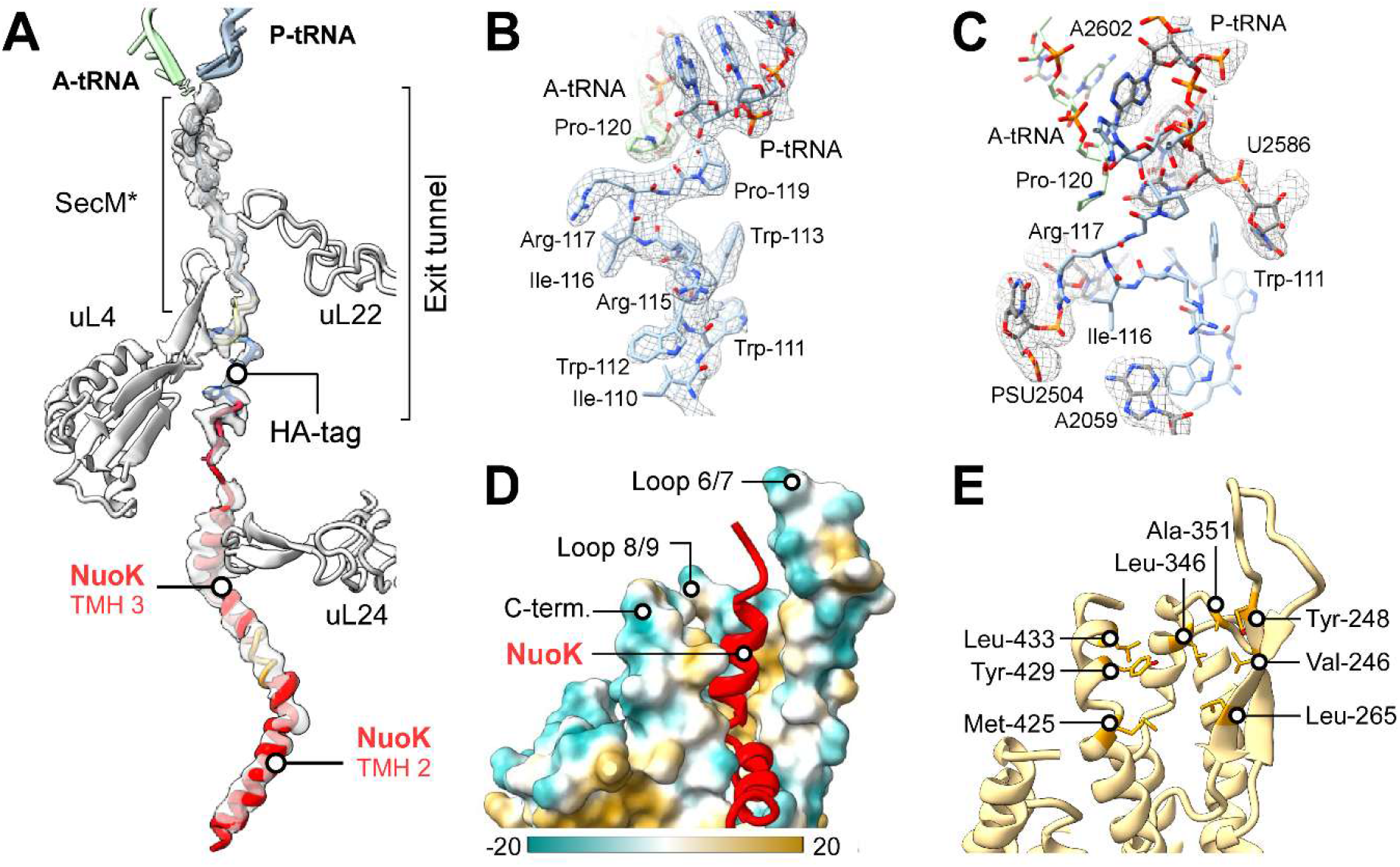
SecYEG and YidC jointly mediate delivery of the nascent chain to the membrane. **(A)** Isolated density of the nascent chain NuoK (in transparent) with fitted model; tRNAs and the ribosomal proteins lining the ribosomal exit tunnel are indicated. **(B)** Isolated cryo-EM density (transparent mesh) of the SecM* stalling sequence in the ribosomal exit tunnel with the fitted model. The residues of SecM* peptide are indicated. **(C)** Model for the SecM*:ribosome interactions. Hallmark bases of the PTC and involved in SecM* binding are shown as in SecM-stalled ribosome (PDB ID 8QOA), fitted into the respective density in (transparent mesh). **(D)** Surface representation of the molecular model for SecY cytoplasmic loops 6/7, 8/9 and the C-terminal tail colored according to the molecular lipophilicity potential (scale bar shown). The emerging NuoK nascent chain (red ribbon) acquires α-helical fold within the hydrophobic pocket. **(F)** The indicated apolar residues form the hydrophobic pocket for the nascent chain folding, as shown in (D).

At the mouth of the exit tunnel, the NuoK nascent chain acquires a helical fold accounting for three C-terminal turns of its TMH 3, which are docked between the SecY loops 6/7 and 8/9, the C-terminal extension of TMH 10 and the tip of the ribosomal uL24 (Fig. 4D). This newly formed arrangement of SecY creates a hydrophobic pocket in the otherwise hydrophilic environment of the exit tunnel and the solvent that allows early folding and accommodation of a short hydrophobic helix (Fig. 4E). As the C-terminal polypeptide of SecY sterically hinders the access to the lateral gate, the emerging nascent chain is not routed into the central channel. Instead, once approaching the cytoplasmic funnel of SecY, NuoK TMH 3 density is kinked by approx. 120° towards the back side of the SecYEG, and so it acquires an interfacial topology laying over TMH 3 of SecE and TMH 10 of SecY (Fig. 5A). These crossed TMHs form a sawhorse-like crevice at the membrane interface oriented toward YidC. In its turn, the paddle domain of YidC covers the nascent NuoK helix from the cytoplasmic side and completes a triangular window for the hydrophobic client. This observation is also consistent with previously proposed interactions of the evolutionary conserved paddle with substrates (McDowell *et al*., 2021). Like the SRP M-domain or Get3/TRC40, the YidC paddle contains a “methionine bristle” for client binding and, indeed, our structure now shows how the membrane-facing Met-408 and Met-409 in CH2 interact with NuoK TMH 3 prior its insertion (Fig. 5A). Notably, replacing these methionine residues with charged residues (Lys, Arg, Asp) or asparagine, but not alanine or serine, rendered a cold-sensitive phenotype in *E. coli* upon depletion of the wild-type YidC (Fig. 5B and Appendix Fig. S8). We speculate that the effect originates from defects in IMP biogenesis, and that the apolar environment provided by the paddle domain is an important factor for the nascent IMP folding at the lipid membrane interface prior the insertion, in agreement with the physical considerations (White & Wimley, 1999).

**Figure 5.**
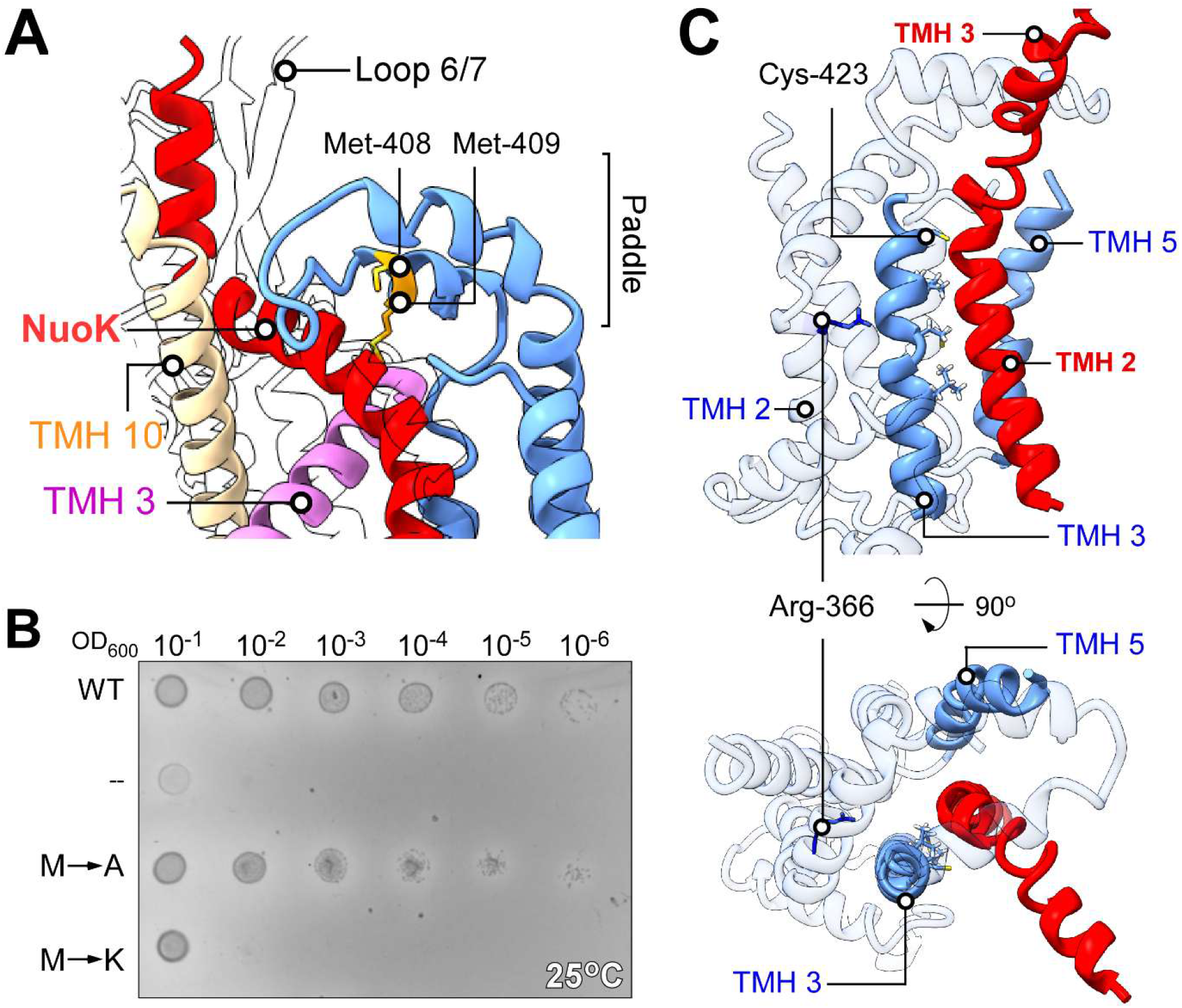
YidC-mediated insertion of the nascent membrane protein. **(A)** The architecture of the nascent chain hand-over interface between SecYEG and YidC. The methionine residues within the YidC paddle domain are indicated. **(B)** Complementation test using wild-type YidC (“WT”), mutants YidC^M408A, M409A^ (“M→A”) and YidC^M408K, M409K^ (“M→K”) and empty vector (“--”) in presence of glucose at 25°C. **(C)** Inserted NuoK TMH 2 is docked at the groove of YidC between TMHs 3 and 5 with the accommodated NuoK TMH 2. The aliphatic residues within YidC TMH 3 of the insertion groove are shown as sticks (Cys-423, Leu-426, Met-430 and Leu-434). Conserved Arg-366 within YidC TMH 2 is indicated. The rest of YidC is shown as semi-transparent ribbon for clarity.

The close SecYEG-YidC association ensures hand-over of the nascent chain to YidC, and the position of NuoK TMH 2 in our structure suggests the insertion route (Fig. 5C). The membrane-inserted NuoK TMH 2 is separated from TMH 3 by a kink, likely a short flexible loop, at the level of YidC Cys-423. NuoK TMH 2 occupies the wide groove between YidC TMHs 3 and 5, with Leu-426, Met-430 and Leu-434 as the closest residues (Fig. 5C), forming the functional site of the insertase described in previous studies (Kedrov *et al*., 2016; Yuan *et al*, 2007). As the groove narrows towards the periplasm, the N-terminal end of NuoK TMH 2 is expelled from the helical bundle of YidC, being fully exposed to the lipid environment. Notably, the visualized NuoK does not reach YidC TMH 2 and does not interact with Arg-366 located deeply in the YidC groove (Fig. 5C). This residue is shown to be important for YidC functionality, as it may destabilize the lipid membrane and ensure polarity and water access within the groove (Chen *et al*., 2017), but also interactions of the conserved arginine with charged/polar residues of client proteins have been reported (Kumazaki *et al*, 2014a; Tsukazaki, 2019). Although not observed in our structure, we cannot exclude that such interactions are formed at other stages of NuoK insertion, or it may be specific for particular nascent IMPs.

### Determinants for the SecYEG-YidC assembly

The C-terminal linker and the stalling peptide of the designed nascent chain retain NuoK^86^-RNC bound to the SecYEG-YidC complex, offering a view on a late insertion intermediate, with NuoK TMH 3 trapped at the interface prior its insertion. In absence of the C-terminal extension, the helix would be released from the ribosome and inserted post-translationally, while the ribosome could detach from SecYEG. To test wether the observed SecYEG-YidC assembly also occurs upon co-translational insertion, we focused on an earlier NuoK intermediate formed prior the nascent chain release (Appendix Fig. S9). The shorter construct NuoK^70^ contained TMH 1 and TMH 2 fully exposed from the ribosome and the N-terminal part of TMH 3 emerging from the tunnel (Fig. 6A and Appendix Fig. S2). The length of the linker/stalling sequence within the ribosomal tunnel matched the length of the replaced C-terminal part of NuoK (30 residues), so the construct approximated the full-length NuoK. Cryo-EM analysis of the assembled NuoK^70^-RNC:SecYEG-YidC complex revealed an architecture very similar to the NuoK^86^-based sample (Fig. 6B and C): While the lateral gate of SecY was closed, YidC was located at the back side of the SecYEG complex, with a slight shift towards the N-terminal half of the translocon, as concluded based on a rigid body fit. Similar to NuoK^86^, a rod-like density accounting for NuoK TMH 3 emerged from the ribosomal tunnel into the hydrophobic pocket formed by the cytoplasmic loops and the C-terminal polypeptide of SecY. Another prominent though less defined density was located above the SecY-SecE crevice, most likely corresponding to the pre-inserted part of NuoK TMH 2. At low contour levels, a density for the membrane-embedded part of NuoK TMH 2 was found near the insertion groove of YidC (Fig. 6B). Despite the limited resolution achieved in presence of NuoK^70^ nascent chain, both reconstructions for the “post-translational” event (NuoK^86^-RNC) and “co-translational” event (NuoK^70^-RNC) reveal nearly identical organization of the assembled SecYEG-YidC machinery. Importantly, YidC was recruited to NuoK^70^ prior TMH 2 was fully embedded into the membrane, thus suggesting that YidC indeed is an insertase rather than a chaperone for the pre-inserted TMHs.

**Figure 6.**
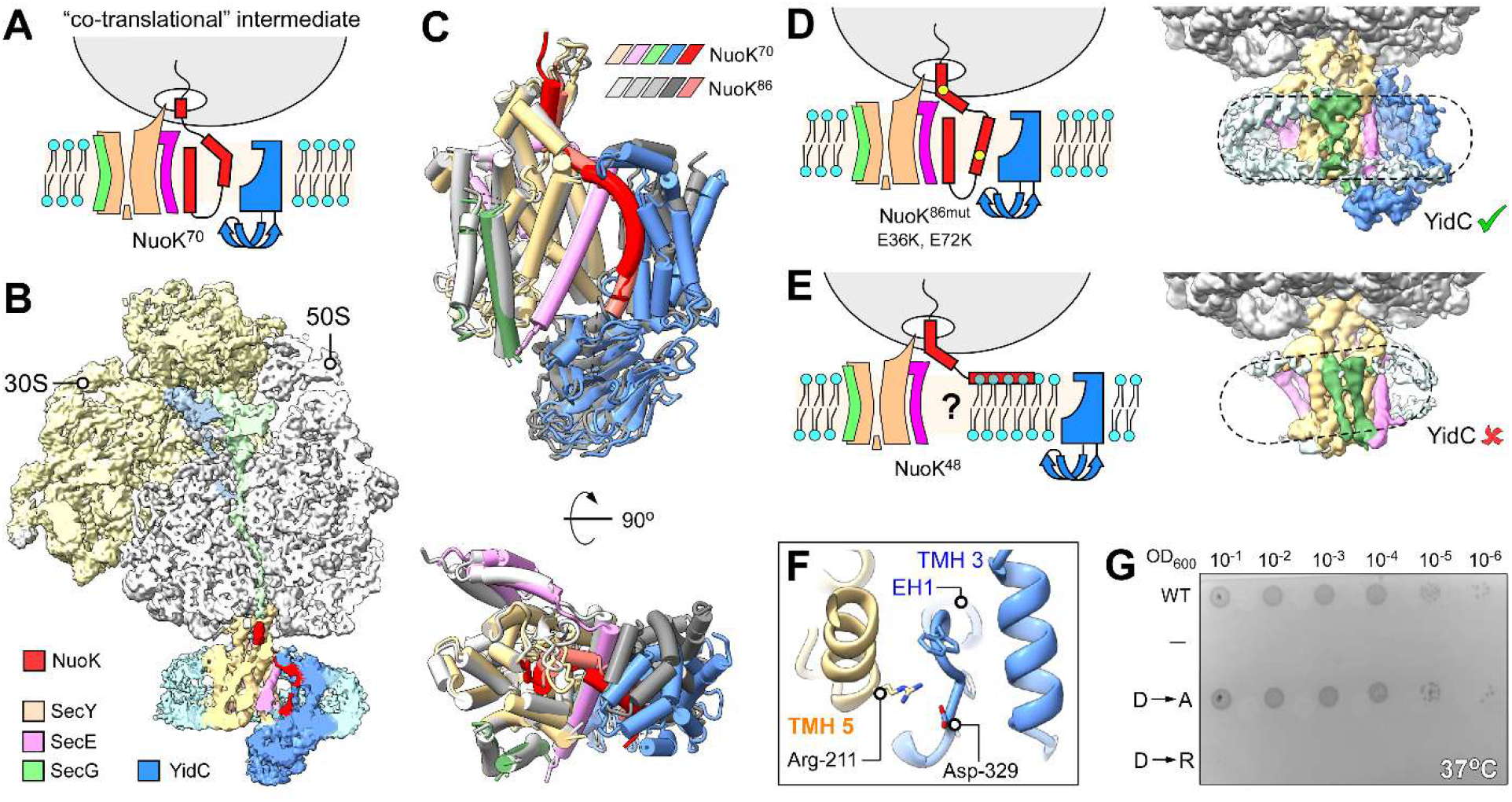
Substrate-induced assembly of the SecYEG-YidC insertase complex. **(A)** Schematic of the NuoK^70^ intermediate mimicking co-translational insertion via the SecYEG-YidC machinery. **(B)** Cryo-EM reconstruction of the NuoK^70^-RNC:SecYEG-YidC assembly (central cross-section) with the individual densities indicated. **(C)** Molecular model of the SecYEG-YidC insertase complex bound to NuoK^70^-RNC. The model of the complex bound to NuoK^86^-RNC is shown in grey. **(D)** Schematic of the NuoK^86^ insertion intermediate bearing E36K, E72K mutations (yellow circles), and the corresponding cryo-EM map (right). The outline of the nanodisc is shown with the dashed line. **(E)** Schematic of the NuoK^48^ insertion intermediate and the corresponding cryo-EM map (right). The outline of the nanodisc is shown with the dashed line. **(F)** View on the molecular model of the SecY-YidC complex focusing on the closest contact site at the periplasmic interface. A putative salt bridge is formed between SecY Arg-211 and YidC Asp-329. **(G)** Complementation test using wild-type YidC (“WT”), mutants YidC^D329A^ (“D→A”) and YidC^D329R^ (“D→R”) and empty vector (“--”) in presence of glucose at 37°C.

We next asked how specific elements of the nascent chain may facilitate recruitment of YidC and determine the complexes’ assembly. As YidC involvement in NuoK biogenesis was previously linked to Glu-36 and Glu-72 residues within TMH 2 and TMH 3 of the nascent IMP, respectively (Price & Driessen, 2010), we substituted both glutamates with lysines and performed cryo-EM analysis of the NuoK^86mut^-RNC:SecYEG-YidC complexes (Fig. 6D; Appendix Fig. S10). Despite the charge inversion and without the glutaraldehyde stabilization, the characteristic YidC density was observed in the same position at the back of SecYEG as upon wild-type NuoK^86^ insertion. Thus, the SecYEG-dependent nascent protein nevertheless promotes recruitment of YidC and the assembly of the insertase complex, though this dynamics could not be addressed in previous studies utilizing protease-protection assays (Price & Driessen, 2010). However, when the length of the nascent chain was reduced to 48 amino acids, so only TMH 1 and TMH 2 of NuoK were exposed from the ribosome (NuoK^48^-RNC), only the RNC-bound SecYEG in its closed state was resolved within the nanodisc (Fig. 6E; Appendix Fig. S11). Notably, neither the C-terminal polypeptide of SecY, nor the helical density exiting the ribosomal tunnel were resolved, suggesting that the the translocon conformation is tuned by the emerging client protein. The signal for YidC was lost, so the insertase remained mobile within the surrounding membrane, and the nanodisc manifested the characteristic tilt by ∼10°. Thus, we concluded that the SecYEG-YidC complex is dynamically assembled to mediate insertion of a nascent IMPs emerging via the “back-of-Sec” route, and the YidC recruitment may not depend on specific charge within the substrate, but may rely on other determinants, such as the length and the folding status of the emerging chain.

Notably, the observed structure of the active SecYEG-YidC complex shows a very similar architecture to the sequence-based model provided by AlphaFold, despite the absence of the ribosome and the nascent chain (Appendix Fig. S5). The direct contacts between SecYEG and YidC are limited to those at the periplasmic side, involving the essential amphipathic helix EH1 of YidC (Fig. 3E) (Kedrov *et al*., 2016; Li *et al*, 2014). Here, SecY Arg-211 (TMH 5) and YidC Asp-329 (end of EH1) potentially form a salt bridge that would stabilize the interaction between both proteins (Fig. 6F and EV4A). Though not conserved through bacterial species, this pair of charged residues is commonly found in γ-proteobacteria, where AlphaFold confidently predicts the SecYEG-YidC association (Appendix Fig. S1). We speculated that disrupting this contact may be detrimental for IMP biogenesis and affect the cell viability. Accordingly, we replaced YidC Asp-329 with either alanine or arginine residues. Both YidC variants could be recombinantly expressed and purified, suggesting that the mutated proteins were correctly folded (Fig. EV4 and C), and immunodetection of the polyhistidine-tagged variants confirmed expression in *E. coli* FTL 10 cells under permissive conditions (Fig. EV4D). When testing the functional complementation of these YidC variants in *E*.*coli* FTL10 strain, we observed that the plasmid-encoded YidC^D329A^ fully supported the cell growth under depletion of the wild-type YidC (Fig. 6G and EV4E). In contrast, introducing a positive charge was detrimental for the cellular viability, similar to the known effect of a minor deletion within the proximate EH1 (Kedrov *et al*., 2016). The effect may be explained by electrostatic repulsion between SecY and YidC, however, we cannot exclude other consequences of the mutation, such as inhibition of YidC-mediated insertion. Future work using a combination of complementary *in vivo* and *in vitro* methods, such as site-specific crosslinking, co-purification, and fluorescence imaging, but also molecular dynamics simulations should provide further insights on the SecYEG-YidC binding interface.

### Comparison of bacterial and eukaryotic insertases

Guiding of the nascent chain to the back side of SecYEG for YidC-mediated insertion manifests an obvious resemblance to the route of multipass IMP biogenesis in the endoplasmic reticulum of eukaryotes (Fig. 7A and B) (Smalinskaite *et al*., 2022). Within the eukaryotic MPT machinery, the ribosome-bound Sec61 translocon builds a core of the substrate-dependent super-complex composed of BOS (“back-of-Sec61”), GEL (“GET- and EMC-like”) and PAT (“proteins associated with the translocon”) complexes (Fig. 7B). We thus compared the arrangement of the *E. coli* SecYEG-YidC with the human insertase machinery trapped with a nascent chain of rhodopsin. The bacterial insertase forms a compact assembly, where YidC is tightly packed against SecYEG and the nascent chain, providing a simple solution to transfer a newly synthesized hydrophobic protein from the ribosome to the membrane-embedded groove of YidC. In contrast, the loosely packed eukaryotic MPT extends far from the Sec61 core and forms a lipid-filled “peninsula” for inserting nascent TMHs. The YidC homolog TMCO1 (a constituent of the GEL complex) is found more than 3 nm away from Sec61 and the nascent TMH, suggesting its role as a membrane chaperone within the MPT rather than an insertase. The position of YidC within the eukaryotic machinery is occupied by TMEM147, a subunit of the BOS complex, that bridges the core translocon with the MPT. Apart from its interactions with Sec61, TMEM147 is known to associate with the ER membrane protein complex (EMC) (Page *et al*, 2024), likely serving as an adaptor subunit in various routes of membrane protein biogenesis.

**Figure 7.**
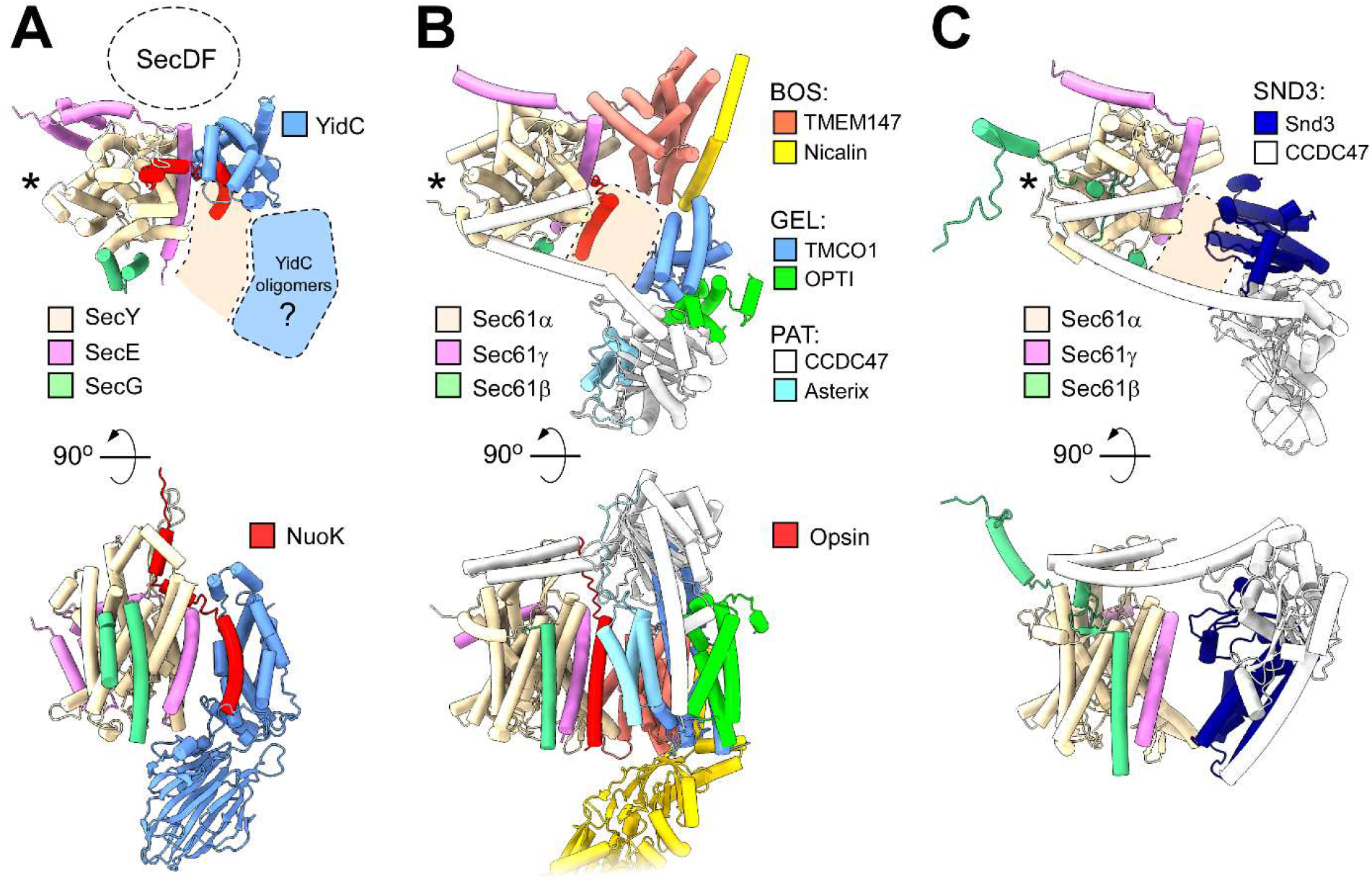
Comparison of the bacterial and eukaryotic IMP insertases. **(A)** Cytoplasmic and side views on *E. coli* SecYEG-YidC assembly and the route of the NuoK nascent chain. The putative position of the accessory SecDF complex is indicated, based on a high-confidence AlphaFold3 prediction (Fig. EV1). A hypothetical position of YidC oligomers is shown in blue. The position of the lateral gate is indicated with an asterisk. **(B)** Cytoplasmic and side views on the eukaryotic multipass membrane protein translocon (MPT) with the subcomplexes and individual subunits indicated (PDB ID 7TUT) The partially resolved route of the opsin nascent chain is indicated. **(C)** Cytoplasmic and side views on the fungal Sec61-SND3 assembly formed in the absence of a client IMP (PDB ID 9I78).

Most recently, a structure of the ribosome-bound Sec61αβγ translocon from the thermophilic fungus *Chaetomium thermophilum* has been visualized in complex with SND3, a membrane machinery involved in biogenesis of a subset of IMPs in eukaryotes (Fig. 7C) (Aviram *et al*, 2016; Yang *et al*, 2025). The observed assembly consists of the Snd3 protein and CCDC47-Asterix, i.e. PAT sub-complex, located at the back of Sec61, thus resembling the architecture of the MPT and SecYEG-YidC. Both for the MPT and SND3, the extended helices of CCDC47 at the cytoplasmic interface reach the N-terminal half of Sec61α and restrict its mobility required for opening the gate, so the nascent IMP is forced to follow the “back-of-Sec” route. In the fungal machinery (Fig. 7C), the entry to the gate is additionally blocked by a cytoplasmic extension of Sec61β, resembling the barrier formed by the C-terminal extension of *E. coli* SecY. Though not evolutionary related to YidC/Oxa1 superfamily, the structure of the Snd3 protein manifests a number of features characteristic for insertases, such as short/tilted TMHs that distort the lipid bilayer, and a polar groove that may serve as the interface for IMP partitioning. While the observed Sec61-SND3 complex likely represents an idle state of the insertase in the absence of a client protein, it certainly highlights the complexity of IMP biogenesis routes beyond the partitioning via the lateral gate.

## Discussion

Variations in TMH topologies and lengths, appearance of partially unfolded regions and charged residues within the membrane core, point to the complexity in IMP biogenesis across the kingdoms of life. Membrane insertion of hydrophobic TMHs via the lateral gate of the universally conserved SecYEG/Sec61αβγ translocon has been described as a general route upon IMP folding. The paradigm has been lately challenged by the discovery of the eukaryotic insertase machinery MPT that facilitates insertion of multipass IMPs at the back of the ribosome-bound Sec61. The discovery of the alternative “back-of-Sec” route in higher eukaryotes (Smalinskaite *et al*., 2022; Sundaram *et al*., 2025), lately substantiated by identifying the ortholog machinery Sec61-SND3 in yeasts (Yang *et al*., 2025), has immediately raised a question about its origins and the evolutionary conservation (Smalinskaite & Hegde, 2023). Here, we show that the minimal bacterial machinery composed of SecYEG and YidC can assemble within the lipid bilayer in the substrate-dependent manner to facilitate insertion of the multipass IMP NuoK via the “back-of-Sec” route. Differently to the lactose permease LacY (Ou *et al*., 2025), the NuoK nascent chain does not pass the lateral gate, but it folds within a hydrophobic pocket formed by SecY loops and leaves via the sawhorse-shaped crevice at the back, where it is encountered by YidC. Notably, folding of a nascent α-helix at the membrane interface appears in excellent agreement with the free energy considerations, as the hydrogen bonds should be established within the helical backbone to enable TMH insertion into the hydrophobic core of the lipid bilayer (White & Wimley, 1999). Once exposed from SecYEG, the pre-folded nascent chain is covered by the partially hydrophobic paddle domain of YidC that assists with handover to the insertion site. The SecYEG-YidC assembly is induced by the nascent chain of sufficient length, and YidC is recruited to assist in insertion/folding of NuoK TMH 2. Although different in the complexity level, the overall architecture of the active SecYEG-YidC insertase manifests a conceptual similarity to the eukaryotic MPT machinery and suggests an evolutionary conservation of the “back-of-Sec” route for IMP biogenesis.

Multiple studies have described interactions of nascent IMPs with the lateral gate of SecYEG and suggested that YidC is located in its proximity to access the emerging nascent TMHs (Ou *et al*., 2025; Petriman *et al*., 2018; Sachelaru *et al*., 2013). In this position, YidC may serve as a chaperone for the newly inserted IMPs, e.g., mediate correct partitioning of moderately hydrophobic helices and ensure assembly of multipass proteins (Fig. 8) (Jauss *et al*., 2019; Petriman *et al*., 2018; Sachelaru *et al*., 2013; Zhu *et al*., 2013). Focused on this route, the previous crosslinking studies extensively investigated the lateral gate region for SecYEG-YidC interactions, while other potential interfaces were barely explored (Fig. EV5). Together with AlphaFold-based modelling, our structural findings demonstrate that an alternative functionally relevant architecture is possible for *E. coli* SecYEG-YidC within the lipid membrane. Notably, the predicted models from other species manifest various, though low-confidence architectures (Appendix Fig. S1). While the insertion pathway is likely to be conserved amongst the organisms, further specie-specific membrane constituents, i.e. accessory proteins and/or the anisotropic and heterogenous lipid environment lacking in the models, may be required to facilitate the assembly of the active insertase.

**Figure 8.**
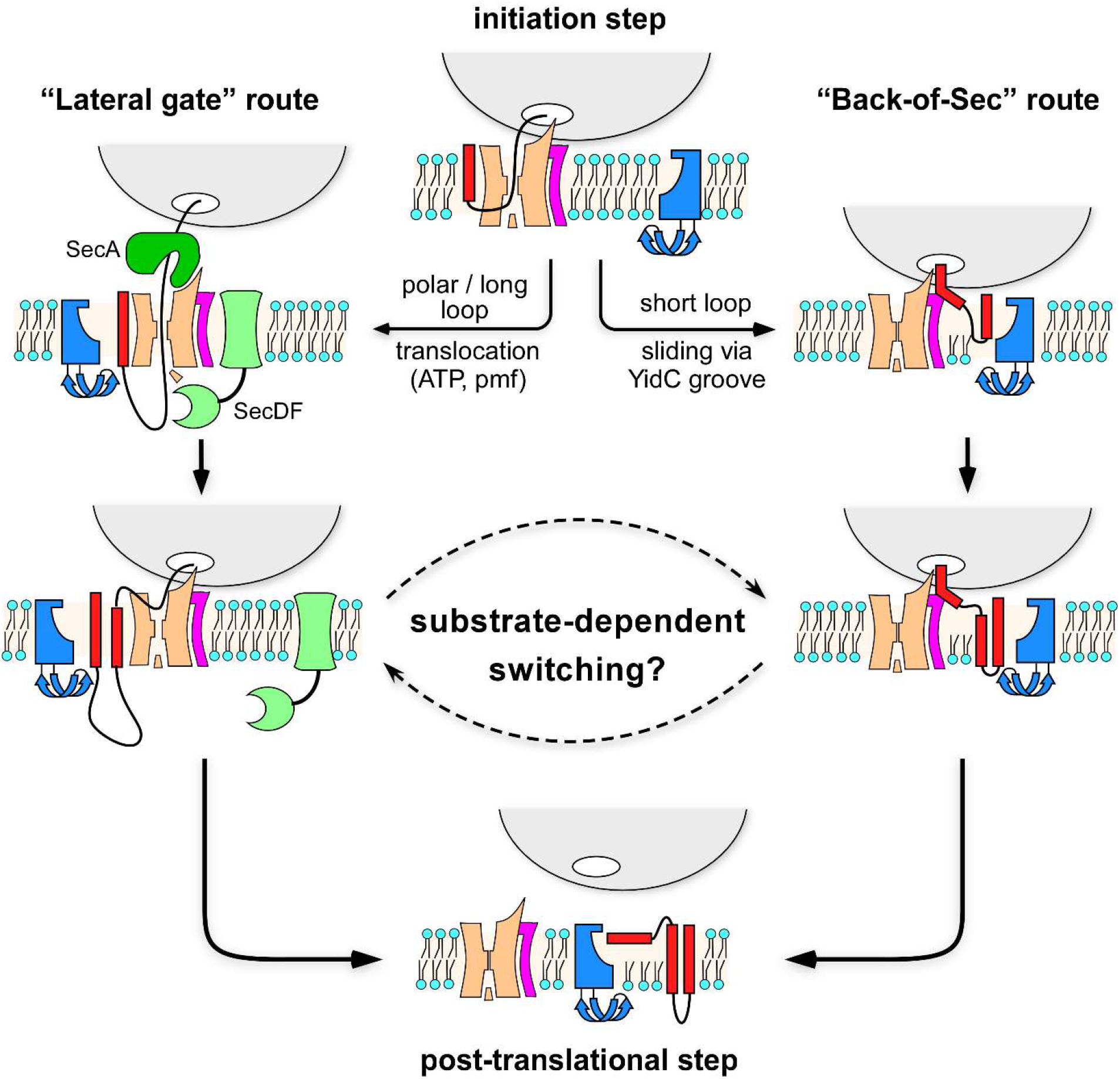
Routes for multipass membrane protein insertion via the bacterial SecYEG-YidC complex. After the SRP-mediated targeting and insertion of the first TMH (here, shown for SecYEG only), membrane partitioning of the following TMHs may occur either via the lateral gate or via the “back-of-Sec” route. The length and the composition of the extramembrane domains determine the biogenesis pathway: SecA/SecDF-mediated translocation of the periplasmic loops requires widening of the lateral gate, and so likely favours the corresponding insertion route. YidC may interact with SecYEG at either of the interfaces to serve as a chaperone, but also assist in insertion of the nascent proteins. Once the translation is completed and the ribosome detaches from SecYEG, insertion of the last TMH occurs post-translationally, being assisted by YidC.

Similar to the eukaryotic MPT (Sundaram *et al*., 2022), assembly of the SecYEG-YidC insertase is dependent on the nascent chain, and the chain composition as well as the propensity to fold may dictate the insertion route. The “back-of-Sec” route in eukaryotes is taken by nascent IMPs lacking long polar/charged loops (Sundaram *et al*., 2025), which otherwise require translocation via Sec61αβγ. The translocation is associated with opening of the translocon and widening of the lateral gate (du Plessis *et al*., 2009), so the primed gate can serve for insertion of a following TMH. Alike, the length and polarity of periplasmic loops dictate the dependence of bacterial IMPs on SecYEG (Soman *et al*, 2014). Biogenesis of a multipass protein, such as LacY, may require multiple translocation events mediated by the ATPase SecA and the proton-powered complex SecDF (Zhu *et al*., 2013), so insertion of the proximate TMHs likely occurs via the lateral gate (Fig. 8) (Ou *et al*., 2025). Importantly, interactions of SecYEG with SecA and a ribosome are mutually exclusive, as both utilize the same binding interface of the translocon (Wu *et al*, 2012). Thus, the ribosome would transiently break the contact with SecYEG upon synthesis of a polar polypeptide stretch, but the nascent chain would serve as a linker to enable re-binding in presence of a following TMH. In contrast, the short loops of NuoK should not depend on SecA-mediated translocation, so the protein manifests the features required for “back-of-Sec” insertion, as observed in the acquired structures. It remains to be explained though, what features of a nascent chain serve as signals for YidC recruitment either at the lateral gate or “back-of-Sec” interface, and whether switching between different routes may occur along insertion of a single IMP.

Notably, the involvement of YidC at the late stage of NuoK biogenesis correlates with the recent findings that both YidC and the evolutionary related EMC complex facilitate insertion of the C-terminal TMHs of polytopic IMPs (Kalinin *et al*, 2024; Wu *et al*, 2024). This final insertion event would occur post-translationally when the translation termination happens soon after the synthesis of the last TMH. NuoK would be an example of such IMP: In absence of the stalling, the nascent polypeptide chain would be released from the peptidyl transferase center, and the membrane insertion of NuoK TMH 3 complex would occur post-translationally. As both YidC and EMC ensure folding of post-translationally delivered clients (Anghel *et al*, 2017; Page *et al*., 2024), the C-terminal TMHs of multipass IMPs may utilize the same recognition and insertion routes, while the SecYEG-YidC assembly would facilitate the efficient substrate delivery (Fig. 8).

The topology acquired by NuoK in the nanodisc is another puzzling outcome of our study. The model based on the resolved TMHs suggests that the N-terminus of the insertion intermediate is oriented into the cytoplasm, thus being inverted in comparison to NuoK within the assembled complex I. The non-native topology could possibly originate from the experimental *in vitro* set-up, e.g. due to the chosen non-native lipid composition, or absence of the physiological electrostatic potential across the membrane, an important factor for the IMP orientation. While NuoK insertion in proteoliposomes efficiently occurred in absence of the membrane potential (Price & Driessen, 2010), the employed protease-protection assay did not account for the resulting protein topology. However, if the electrostatic potential was a decisive factor, we would expect to find NuoK in two different topologies in cryo-EM reconstructions, possibly utilizing different insertion routes, while only one class was experimentally observed. As the observed topology is likely determined by the insertion of NuoK TMH 1, further analysis should assess this early event. However, another scenario should be considered upon the interpretation: Several IMPs, such as four-TMH EmrE, were shown to undergo partial or even complete topology inversion along their folding pathway, and a single charged residue here can be decisive for the final orientation (Lu *et al*, 2000; Seppala *et al*, 2010; Seurig *et al*, 2019; Woodall *et al*, 2017). As NuoK does not contain long loops between the TMHs but contains a stretch of three arginine residues within its C-terminal end, the protein may acquire its correct topology in the cellular membrane once the complete polypeptide chain is released from the ribosome. The flip-flop of the inserted TMHs may be promoted by the proximate YidC, as the insertase would distort the lipid bilayer and so destabilize the NuoK intermediate that eventually results in the new topology, in accordance to a model suggested for the eukaryotic MPT (Smalinskaite & Hegde, 2023).

Our study delivers a new view on the organization of the bacterial SecYEG-YidC insertase machinery, and the revealed architecture is calling for detailed investigation on its dynamics, as well as re-evaluation of the earlier data. One current limitation of the employed nanodisc-based system is the pre-defined stoichiometry of SecYEG and YidC within the complex, while YidC is substantially more abundant in the bacterial membrane (Urbanus *et al*, 2002). Thus, even larger insertase complexes may be envisioned for the cellular membrane, where multiple YidC copies could manage biogenesis of polytopic IMPs (Fig. 7). Experiments on the native bacterial membranes, such as those by super-resolution fluorescent imaging and cryogenic electron tomography (Gemmer *et al*, 2023; Wirth *et al*, 2023), as well as reconstituted systems will be of the utmost importance to explore such modes of interaction, but also to determine exact roles played by YidC along insertion of client IMPs, and so deliver detailed and possibly direct insights on the key process of membrane protein biogenesis.

## Methods

### Molecular cloning

The primers were synthesized by Eurofins Genomics and Merck/Sigma-Aldrich. The used restriction enzymes and cloning kits, incl. the Phusion High-Fidelity DNA polymerase and the Gibson assembly kits were purchased by New England Biolabs. Sequencing was conducted by Eurofins Genomics and Microsynth AG.

The genes encoding for SecYEG-YidC complex were cloned in a pBAD-TOPO (Thermo Fisher Scientific) derived vector. The gene encoded for *E. coli* YidC (1-535) was fused to the N-terminus of SecE with a sequence coding for HRV-3C protease cleavage site inserted in between; the construct also contained a N-terminally decahistidine-tagged SecY and unmodified SecG. The construct was created using conventional restriction-ligation cloning techniques resulting in the plasmid pEM472. To study the effects of point mutations within YidC in complementation assays, the YidC mutants were cloned into pTrc99A-based plasmid pKAD107 (Kedrov *et al*., 2013). For the expression of the YidC mutants, the corresponding genes were cloned into pBAD-based plasmid (Kedrov *et al*., 2016).

For the IMP substrate, the DNA encoding 86 amino acids of *E. coli* NuoK and the C-terminal extension containing the HA-tag for immunodetection, a linker (`GGSG`) and the SecM* stalling sequence (FSTPVWIWWWPRIRGPP) (Gersteuer *et al*., 2024) was synthesized (GenScript Biotech, Netherlands), and the construct was cloned into pRSET vector (Thermo Fisher Scientific) via SalI/HindIII restriction sites. The genes encoding shorter nascent chains were prepared by removing fragments of the initial plasmid corresponding to NuoK residues 71-86 for NuoK^70^ and residues 49-86 for NuoK^48^ via PCR, followed by blunt-end ligation. For the analysis *in vivo*, the sequences the genes encoding for the nascent chains were amplified by PCR, while introducing NheI and HindIII restriction sites, and the resulting PCR products were ligated into pET24a vector.

### Isolation of ribosome-associated complexes from E. coli membranes

*E. coli* BL21(DE3) cells were transformed with the pET24a-based plasmids encoding for NuoK intermediates with SecM* stalling sequences, or empty pET24a vector. The cells were grown in LB medium supplemented with 50 µg/mL kanamycin at 37°C while shaking at 180 rpm. Upon reaching OD_600_ 0.8, protein expression was induced by 0.5 mM IPTG and carried out for 30 min. Cells were harvested, lysed using a microfluidizer (M-110P, Microfluidics Corp.), and the debris was removed by centrifugation at 18000 g for 10 min (SS34 rotor, Sorvall/Thermo Scientific). Membranes were isolated by ultracentrifugation at 42000 rpm for 1 h (45 Ti rotor, Beckman Coulter). Membranes were then suspended in 50 mM Hepes-KOH pH 7.4, 150 mM KOAc, 25 mM Mg(OAc)_2_, 5 % glycerol, 200 µM tris(2-carboxyethyl) phosphine (TCEP), 1 % n-dodecyl β-maltoside (DDM; Glycon Biochemicals GmbH) for 1 h at 4° C. The solubilized material was loaded on sucrose cushion (50 mM Hepes-KOH pH 7.4, 150 Mm KOAc, 25 mM Mg(OAc)_2_, 0.05% DDM, 200 µM tris(2-carboxyethyl) phosphine (TCEP), 1.1 M sucrose) and centrifuged for 1 h at 110.000 g (AT3 rotor, Sorvall/Thermo Scientific). The supernatant was carefully removed, and the pellet resuspended in 50 mM Hepes-KOH pH 7.4, 150 mM KOAc, 25 mM Mg(OAc)_2_, 5 % glycerol, 200 µM tris(2-carboxyethyl) phosphine (TCEP), 0.05 % DDM prior to loading on SDS-PAGE and subsequent Western blotting.

After SDS–PAGE and transfer onto a methanol-activated PVDF membrane, the membrane was blocked for 2 h at 20 °C in TBS-T buffer supplemented with 5 % (w/v) milk powder. To detect YidC, polyclonal rabbit antibodies were raised against the YidC periplasmic domain (residues 26 to 331; Davids Biotechnologie GmbH). The serum containing the antibodies was diluted 1:1000 in TBS-T supplemented with 2 % (w/v) bovine serum albumin (BSA), and incubated for 2 h at 20 °C. For detection, horseradish peroxidase (HRP)-conjugated anti-rabbit secondary antibodies was applied at 1:10,000 dilution in TBS-T complemented with 2 % (w/v) BSA and incubated for an additional 2 h at 20 °C. Detection reaction was performed with the Clarity Western ECL substrate (Bio-Rad Laboratories) and imaged with the Amersham Imager 680 (GE Life Sciences).

### Expression of the membrane insertases

*E*.*coli* C41(DE3)Δ*ompF-*Δ*acrAB* strain (Kanonenberg *et al*, 2019) transformed with the pEM472 plasmid encoding for the SecYEG-YidC fusion construct was grown in LB medium supplemented with 100 µg/mL ampicillin at 37°C while shaking at 180 rpm. Upon reaching OD_600_ 0.6, expression of the SecYEG-YidC complex was induced by adding 0.5 % arabinose (w/v) and carried out for 2.5 h. Cells were harvested and lysed using a microfluidizer (M-110P, Microfluidics Corp.), and the debris was removed by centrifugation at 18000 g for 10 min (SS34 rotor, Sorvall/Thermo Scientific). Membranes were isolated by ultracentrifugation at 42000 rpm for 1 h (45Ti rotor, Beckman Coulter). Membranes were then suspended in 50 mM Hepes-KOH pH 7.4, 150 mM KOAc, 5 % glycerol, 200 μM tris(2-carboxyethyl)phosphine (TCEP), 1 % DDM for 1 h at 4°C. The solubilized material was centrifuged at 21,380 x *g* for 10 min at a tabletop centrifuge (Hermle Z 216 MK, Hermle Labortechnik GmbH) and the supernatant was incubated with Ni^2+^-NTA agarose resin (Macherey-Nagel GmbH & Co. KG) for 1 h at 4°C. The resin was washed with 50 mM Hepes-KOH pH 7.4, 500 mM KOAc, 5 % glycerol (v/v), 200 μM TCEP, 0.05 % DDM, 10 mM imidazole and the target protein was eluted with 50 mM Hepes-KOH pH 7.4, 150 mM KOAc, 5 % glycerol, 200 μM TCEP, 0.05 % DDM, 300 mM imidazole. The elution fractions were concentrated and subjected to size exclusion chromatography (SEC) in 50 mM Hepes-KOH pH 7.4, 150 mM KOAc, 5 % glycerol, 0.05% DDM using Superdex 200 Increase GL 10/300 column with ÄKTA Pure set-up (Cytiva). The protein concentration was determined spectrophotometrically (NeoDot, NeoBiotech) using the calculated extinction coefficient of 134,000 M^-1^ cm^-1^. Expression of YidC mutants, YidC^D329A^ and YidC^D329R^, was achieved using pBAD-based plasmid and the proteins were solubilized and purified in DDM according to the previously published protocol (Kedrov *et al*., 2016).

### Assembly of SecYEG-YidC nanodiscs

To prepare the nanodiscs, liposomes were first formed using 1-palmitoyl-2-oleoyl-glycero-3-phosphocholine (POPC; 70 mol %) and 1-palmitoyl-2-oleoyl-sn-glycero-3-phospho-(1-rac-glycerol) (POPG; 30 mol %) (Avanti Polar Lipids, Inc). POPC was chosen as a zwitterionic component to substitute phosphatidylethanolamine (PE) naturally occurring in *E. coli* membranes. Due to its cylindrical shape, POPC readily forms planar membranes, and so ensures higher homogeneity among formed nanodiscs. Furthermore, both POPC and POPG manifest order-disorder transition below 0°C, in contrast to POPE (transition temperature at 25°C), so the composite lipid bilayer should remain in the fluid state along nanodiscs formation. The lipids were mixed from chloroform stocks to achieve the desired ratio, the solvent was evaporated and the liposomes were prepared, as described (Kater *et al*., 2019). The detergent-purified SecYEG-YidC complexes were reconstituted in MSP2N2-based nanodiscs in a protein:MSP:lipid molar ratio of 1:4:400 following the published protocol (Kater *et al*., 2019). After forming the nanodiscs, the linker connecting YidC and SecE was cleaved by HRV-3C protease for 1 h at 4°C and SEC was performed in 50 mM HEPES-KOH pH 7.4, 150 mM KOAc, 5 % glycerol using Superose 6 Increase GL 10/300 column with ÄKTA Pure set-up. SEC fractions containing SecYEG-YidC nanodiscs were concentrated to ∼10 µM using Amicon Ultra-4, Ultracel 30 K centrifugal filters (Merck/Millipore). Mass photometry measurements were performed using Two^MP^ instrument (Refeyn Ltd.) calibrated using β-amylase present in various oligomeric states, with the molecular masses ranging from 56 kDa (monomer) to 224 kDa (tetramer).

### Cell-free protein synthesis

*E. coli* S30 extract for CFPS was prepared based on previously published protocols (Zubay, 1973). Briefly, *E*.*coli* BL21(DE3) cells were transformed with TargoTron pAR1219 plasmid (Sigma-Aldrich) encoding for T7 RNA polymerase. 2 L of 2x YPTG media was inoculated with 100 mL overnight cultures, the cells were grown to OD_600_ 0.5, and the T7 RNA polymerase expression was induced with 1 mM IPTG. The cells were further grown to reach OD_600_ 1.0 and then they were harvested at 7500 rpm for 15 min (FiberLite F8-6 × 1000y rotor, Piramoon Technologies Inc.). The cell pellet was washed three times with 10 mM Tris-acetate pH 8, 60 mM KOAc, 14 mM Mg(OAc)_2_, and 1 mM PMSF, and the pellet was resuspended in the same buffer at the ratio of 1 mL per 1 g pellet. Subsequently, cells were lysed by sonication (10 times, 15 s on, 30 s off, 50 % power, 5 pulsed cycles) (Sonopuls GM2200, Bandelin). The lysate was cleared via two-steps centrifugation, 12,000 x *g* for 15 min and 30,000 x *g* for 30 min (S120 AT6 rotor, Sorvall /Thermo Fisher). The supernatant was aliquoted and stored at -75°C.

The CFPS reaction was composed of 40 % S30 extract, the master mix (10 mM ammonium acetate, 130 mM KOAc, 33 mM sodium pyruvate, 1.5 mM spermidine, 1 mM putrescine, 4 mM sodium oxalate, 1.2 mM ATP, 0.85 mM of GTP, CTP and UTP, 34 μg/ml folinic acid, 170.6 μg/mL of *E*.*coli* tRNA MRE 600 (Roche Diagnostics GmbH), 0.33 mM NAD^+^, 0.26 mM coenzyme A and 2 mM of each amino acid), and Mg(OAc)_2_. The optimal concentration of Mg(OAc)_2_ was identified for each new batch of the S30 extract upon screening within the range of 0 to 12 mM, while using synthesis of the yellow fluorescent protein as a read-out. For NuoK synthesis, at least 7 ng/μL plasmid DNA encoding the nascent chain was added to the reaction, as well as at least 100 nM of the nanodisc-reconstituted SecYEG-YidC. CFPS reactions were performed at 37°C for 1 h while shaking at 450 rpm. The synthesis/stalling was evaluated via Western-blotting using monoclonal antibodies against the HA-tag (sc-7392, Santa Cruz Biotechnology).

### Isolation of ribosomes from CFPS reactions

To isolate the ribosomes, 10-40 % linear sucrose gradients were formed in SW40 tubes (Beckman Coulter) using the Gradient Station (BioComp Instruments). CFPS reactions (100 μL) were loaded on top of the gradients and the samples were centrifuged for 16 h at 16,500 rpm (SW40 Ti rotor; Beckman Coulter). The gradients were fractionated using the Gradient Station while monitoring the absorbance at 280 nm (A_280_). The peaks with the ribosomal fractions occurring in the sucrose concentration range of 20-25 % were pooled together and incubated with Ni^2+^-NTA agarose resin (Macherey-Nagel GmbH & Co. KG) for 1 h at 4°C. The resin was washed with 50 mM Hepes-KOH pH 7.4, 500 mM KOAc, 25mM Mg(OAc)_2_, 10 mM imidazole. The complexes were eluted with 50 mM Hepes-KOH pH 7.4, 150 mM KOAc, 25 mM Mg(OAc)_2_ supplemented with 300 mM imidazole, and the eluate was concentrated using Amicon Ultra-4 Ultracel 30 K centrifugal filters (Merck/Millipore) while exchanging the buffer to 50 mM Hepes-KOH pH 7.4, 150 mM KOAc, 25 mM Mg(OAc)_2_. The presence of the ribosome-bound nascent chain was confirmed by immunoblotting against the HA-tag. The concentration of ribosomes was then estimated by measuring the absorbance at 260 nm (A_260_).

In order to improve the resolution of the cryo-EM data on NuoK^86^-RNC:SecYEG-YidC and NuoK^70^-RNC:SecYEG-YidC, glutaraldehyde was added to the purified samples. The sample was diluted 5-fold to prevent inter-particle crosslinking, and it was then incubated with 0.1 % glutaraldehyde (v/v) for 15 min on ice and quenched by 100 mM Tris-HCl pH 7.5. Afterwards, the sample was concentrated using Amicon Ultra-4 Ultracel 30 K filters, flash-frozen and stored at -75°C before grid preparation.

*Cryo-EM sample preparation and data collection* Isolated NuoK-RNC:SecYEG-YidC samples were supplemented with (1H, 1H, 2H, 2H-perfluorooctyl)-β-D-maltopyranoside (FOM, Anatrace) to a final concentration of 0.03% to favor random orientation of the particles and plunge frozen. For each grid, 3.5 μL of the sample was applied onto glow-discharged Quantifoil Cu 300 mesh R3/3 grids with an additional 2 nm layer of carbon. After a waiting time of 45 s, the grids were blotted for 3 s and plunge frozen in liquid ethane at 4°C and 100% humidity using a Vitrobot Mark IV (Thermo Fisher Scientific). Data collection for RNC-NuoK:SecYEG-YidC samples was performed at 300 keV using a Titan Krios microscope equipped with a Falcon 4i direct electron detector and a SelectrisX imaging filter (all Thermo Fisher Scientific) at a pixel size of 0.727 Â. Dose-fractioned movies were collected in a defocus range from -0.5 to 3.5 μm and with a total dose of 60 e^-^ per Å^2^, fractionated in 60 frames to obtain a total dose of 1 e^-^ per Å^2^ per frame. Gain correction, movie alignment and summation of movie frames were performed using MotionCor2 (Zheng *et al*, 2017). Further processing, including CTF estimation, was carried out in cryoSPARC v4.4 (Punjani *et al*, 2017).

### Data processing

For the crosslinked NuoK^86^-RNC:SecYEG-YidC complex, 27,660 micrographs were selected. Blob Picker was used to pick 1,420,623 particles which were sorted by 2D classification, yielding a subset of 636,995 particles. An *ab-initio* job with 2 classes was run to further clean the particle set. A consensus refinement of 582,663 particles resulted in a map of 70S with clear extra density below the exit tunnel. A soft mask covering this region (accounting for nanodisc, SecYEG-YidC and eventually the NuoK nascent chain) was used to sort the particles into 6 classes, using a 3D Classification job. Two of the classes, representing a total of 221,144 particles, displayed a strong SecYEG and YidC density. The class with best resolved nascent chain density containing 113,368 particles (19.5 %) was selected and refined to a final resolution of 2.44 Å. Local refinement was performed on the tunnel exit region yielding a map with a final resolution of 3.76 Å. This map was used to build the atomic model for the NuoK:SecYEG-YidC assembly (PDB: 9RBF; EMDB:53892; EMDB:53893 (local refinement); EMDB:53894 (composite map)). Data processing for this dataset is summarized in Fig. EV2.

For the non-crosslinked NuoK^86^-RNC:SecYEG-YidC assembly, 10,234 micrographs were selected and manually curated. A total of 693,102 particles was picked with Blob Picker and sorted by 2D classification to generate templates for the Template Picker job. From 2,300,222 template-picked particles, 406,978 were selected after 2D classification. An ab-initio job (three classes) allowed further cleaning of the data set. A consensus refinement with a set of 217,418 particles resulted in a map of 70S ribosome with clear extra density below the exit tunnel. Focused sorting below the exit tunnel region into five classes yielded a class of 33,912 particles showing density for the extramembrane P1 domain of YidC. These particles were used to train a TOPAZ picking model, which resulted in 360,610 particles. The particles were further curated and focused sorted into a final class of 70,670 particles. This class was refined to a final resolution of 2.73 Å (EMD-53587). A local refinement in the SecYEG-YidC region yielded a map with a final resolution of 3.19 Å (EMD-53589). Data processing for this dataset is summarized in Appendix Fig. S6.

For the NuoK^70^-RNC:SecYEG-YidC complex, an initial set of 1,145,565 particles was picked using Blob Picker based on 31,572 micrographs. A final set of 956,488 particles was selected after 2D classification for homogeneous refinement, using an initial map of the 70S ribosomes obtained by an *ab-initio* reconstruction with two classes. Next, 3D classification was performed using a mask for the nanodisc region, resulting in two classes that differed by the presence of YidC. The SecYEG-YidC class (529,825 particles) was further sorted into three classes using a 3D classification job with a mask in the nanodisc region. This resulted in a class of 113,106 particles (21.7%) with well-resolved YidC features. Using a mask around YidC, those particles were further sub-sorted into a class containing only SecYEG and a class representing the SecYEG-YidC assembly (58,063 particles). The SecYEG-YidC class was refined yielding a final map with 2.85 Å resolution. Local refinement with a mask for the nanodisc region yielded a map with a final resolution of 2.90 Å. These maps were used to build an atomic model for the NuoK^70^-RNC:SecYEG-YidC complex (PDB: 9T5X; EMDB:55570; EMDB:55571 (local refinement); EMDB:55598 (composite map)). Data processing for this dataset is summarized in Appendix Fig. S9.

For the NuoK^86mut^-RNC:SecYEG-YidC complex assembled upon NuoK^E36K, E72K^ synthesis Blob Picker was used to pick an initial set of 586,721 particles from 9,099 micrographs. The particles were curated and after 2D classification, a subset of 287,399 particles was used to generate an *ab-initio* reconstruction (two classes) of 70S ribosomes with extra density for the insertase complex. The same set was further used to generate templates for template picking, which resulted in 2,057,403 particles, from which a final set of 56,980 particles was obtained after extensive 2D and 3D classification. The particles were fed into a TOPAZ training job, obtaining 433,249 particles after picking and extraction. These particles were used to generate a consensus refinement and focused sorted into eight classes. The class with the best resolved insertase region, comprising 48,952 particles, was further refined to generate a reconstruction with final resolution of 2.87 Å (EMD-53584). A local refinement with a mask in the translocon region generated a map of 3.20 Å (EMD-53585). Data processing for this dataset is summarized in Appendix Fig. S10.

For the NuoK^48^-RNC:SecYEG-YidC complex assembled upon synthesis of the early NuoK intermediate, an initial set of 741,311 particles was picked using a Blob Picker job based on 21,440 micrographs. The particles were cleaned by 2D classification jobs to yield a final 511,016 particle set. An *ab-initio* reconstruction (two classes) job was used to generate an initial map of the ribosome-bound SecYEG in the nanodisc. Successive, focused 3D-classifications from a consensus refinement were used to obtain a final class of 66,486 particles. These particles were refined to a final resolution of 2.62 Å (EMD-53560). A local refinement with a mask in the translocon region resulted in a reconstruction of 2.90 Å (EMD-53568). Data processing for this dataset is summarized in Appendix Fig. S11.

For visualization of all cryo-EM maps, consensus maps were generated consisting of isolated density for the 70S ribosome from the global refinements and isolated density for the nanodisc-embedded insertase from the local refinements. Statistics to cryo-EM data collection and model refinement is provided in Appendix Table S2.

### Model building and refinement

The molecular model for the crosslinked NuoK^86^-RNC:SecYEG-YidC was built as follows: For the 70S ribosome a previously released model based of high-resolution cryo-EM maps of a 70S ribosome and a SecM-stalled RNC were used as templates (PDB IDs 7K00, 8QOA) (Gersteuer *et al*., 2024; Watson *et al*, 2020). The model for the tRNA-Gly (in SecM) was changed to tRNA-Pro and the mRNA model was adjusted from GCU-GGC-CCU (Ala-Gly-Pro in SecM) to GGU-CCU-CCG (Gly-Pro-Pro in SecM*). A model for the SecYEG-YidC assembly was generated based on the rigid body fitting the AlphaFold3 prediction of this complex (see also Appendix Fig. S4) that was fitted into the locally refined cryo-EM density with only minor adjustments (see also Fig. EV2). For the nascent chain, the modified SecM* sequence could be modelled *de novo* based on well-resolved density for this region. Less-resolved density was present to fit the backbone and a few bulky side chains for major parts of glycine-serine linker and the HA-tag. NuoK TMHs 2 and 3 were identified based on the rod-like shape of extra density present at the mouth of the exit tunnel and between SecY and YidC. While at the given resolution we cannot be sure about the exact register, we positioned residues 73-82 of NuoK THM 3 into the corresponding density supporting the predicted α-helical conformation. 70S ribosome and SecYEG-YidC atomic models were processed independently in multiple rounds of manual real-space refinement in Coot v0.9.8.95 (Emsley & Cowtan, 2004). The models were later merged and further refined in Phenix v1.20.1-4487 (Adams *et al*, 2010). The NuoK^86^-RNC:SecYEG-YidC model was then used as a template to build a model for the crosslinked NuoK^86^-RNC:SecYEG and the crosslinked NuoK^70^-RNC:SecYEG-YidC structures. For the former, the YidC model as well as the section of NuoK not visible in the map was deleted from the template, for the latter only the NuoK sequence was adjusted accordingly. Both models were real-space refined in Coot and Phenix. The Molprobity tool was used for model validation. Visualization was done in UCSF ChimeraX v1.9 (Goddard *et al*, 2018).

### Complementation assay

The complementation assay was prepared as described before (Kedrov *et al*., 2013). Briefly, a single colony of FTL10 cells (Hatzixanthis *et al*, 2003) transformed with either YidC-encoding plasmid or empty pTrc99A vector was grown in LB medium with 0.2% arabinose, 25 µg/mL kanamycin and 100 µg/mL ampicillin for 16 h at 37°C at 180 rpm. The overnight cultures were diluted to OD_600_ of 0.05 and grown until the early logarithmic phase before diluting them all to OD_600_ of 0.1 and doing a subsequent serial dilution. 5 µL of each diluted culture were transferred on the plates (LB agar supplemented with 25 µg/mL kanamycin, 100 µg/mL ampicillin and either 0.2% arabinose or 0.2% glucose). The plates were incubated at 37°C for 16 h or at 25°C for 24 h and photographed with the Amersham Imager 680 (GE Life Sciences). For visualization purposes, brightness levels were adjusted with GIMP 2.10 software.

To test the expression of YidC mutants in *E. coli* FTL 10 strain under permissive conditions, the cells were transformed with pTrc99A-based plasmids encoding for His-tagged YidC variants (Kedrov *et al*., 2013) and grown overnight at 37 °C in LB medium supplemented with 25 µg/mL kanamycin, 100 µg/mL ampicilline and 0.2% arabinose. The cultures were adjusted to the same optical density, and the cells were lysed in SDS-PAGE sample buffer. Subsequent anti-His Western blot analysis (primary antibodies DIA-900-200, Dianova) confirmed expression of all tested YidC^His^ variants.

## Data Availability

The cryo-EM structural data have been deposited in the Electron Microscopy Data Base (EMDB) and the Protein Data Bank (PDB) repositories under the accession numbers EMD-53892, EMD-53893, EMD-53894 and PDB-9RBF (NuoK^86^-RNC:SecYEG-YidC (crosslinked)); EMD-55570, EMD-55571, EMD-55598 and PDB-9T5X (NuoK^70^-RNC:SecYEG-YidC (crosslinked)); EMD-53587 and EMD-53589 (NuoK_86_-RNC:SecYEG-YidC); EMD-53584 and EMD-53585 (NuoK_86mut_-RNC:SecYEG-YidC); EMD-53560 and EMD-53568 (NuoK_48_-RNC:SecYEG-YidC).

## Acknowledgements

The authors would like to thank Hanna Kratzat, Susanne Rieder and Charlotte Ungewickel for the support with the cryo-EM collection and initial data analysis, Joanna Musial for assistance with cloning and biochemical characterization of the SecYEG-YidC construct, Laura Czech and Gert Bange for the support upon the development of CFPS procedure, and Arnold J.M. Driessen and Florian Altegoer for discussions and valuable advices. The work was funded by German Research Foundation (DFG, grants Ke1879/3 and Collaborative Research Center 1208, project A10 to A.K.) and European Research Council (Advanced Grant “CryoTranslation” to R.B.)

## Expanded View Figures

**Figure EV1.**
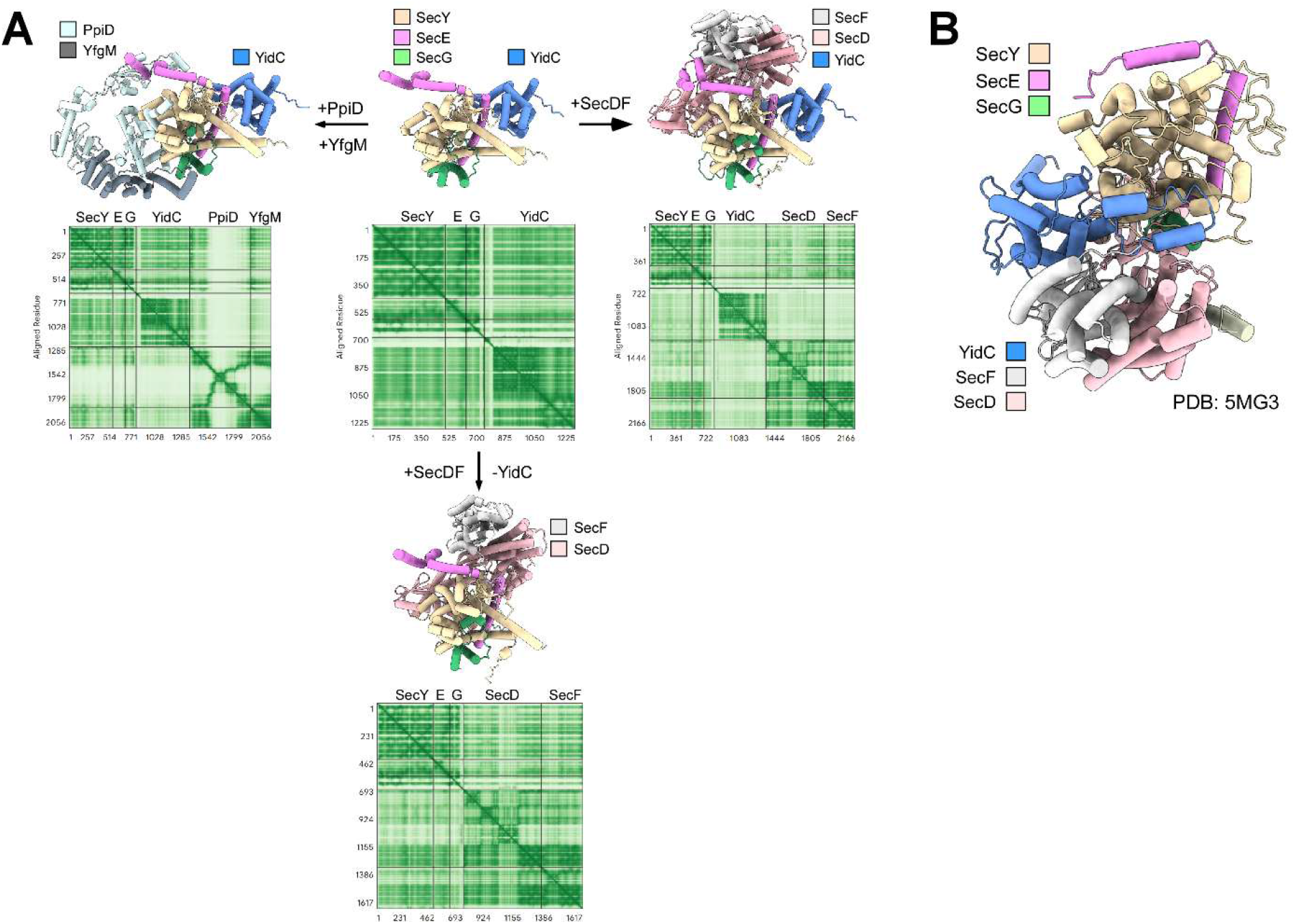
Potential interaction network of *E. coli* SecYEG. **(A)** AlphaFold3-based models and PAE plots of putative complexes formed by *E. coli* SecYEG with the accessory proteins. YidC is docked at the back of SecYEG with high confidence (low PAE levels), though co-assembly with SecDF negatively affects the model confidence for both YidC and SecDF. This may be explained by steric hindrance and partial overlap of the binding sites at SecYEG. Single-span protein YajC manifested random positions between the models, with high PAE scores, so it was excluded from analysis. The generated AlphaFold models are provided in the Dataset EV1. **(B)** Model of the SecYEG-YidC-SecDF assembly in detergent, based on the low-resolution cryo-EM reconstruction (PDB ID 5MG3).

**Figure EV2.**
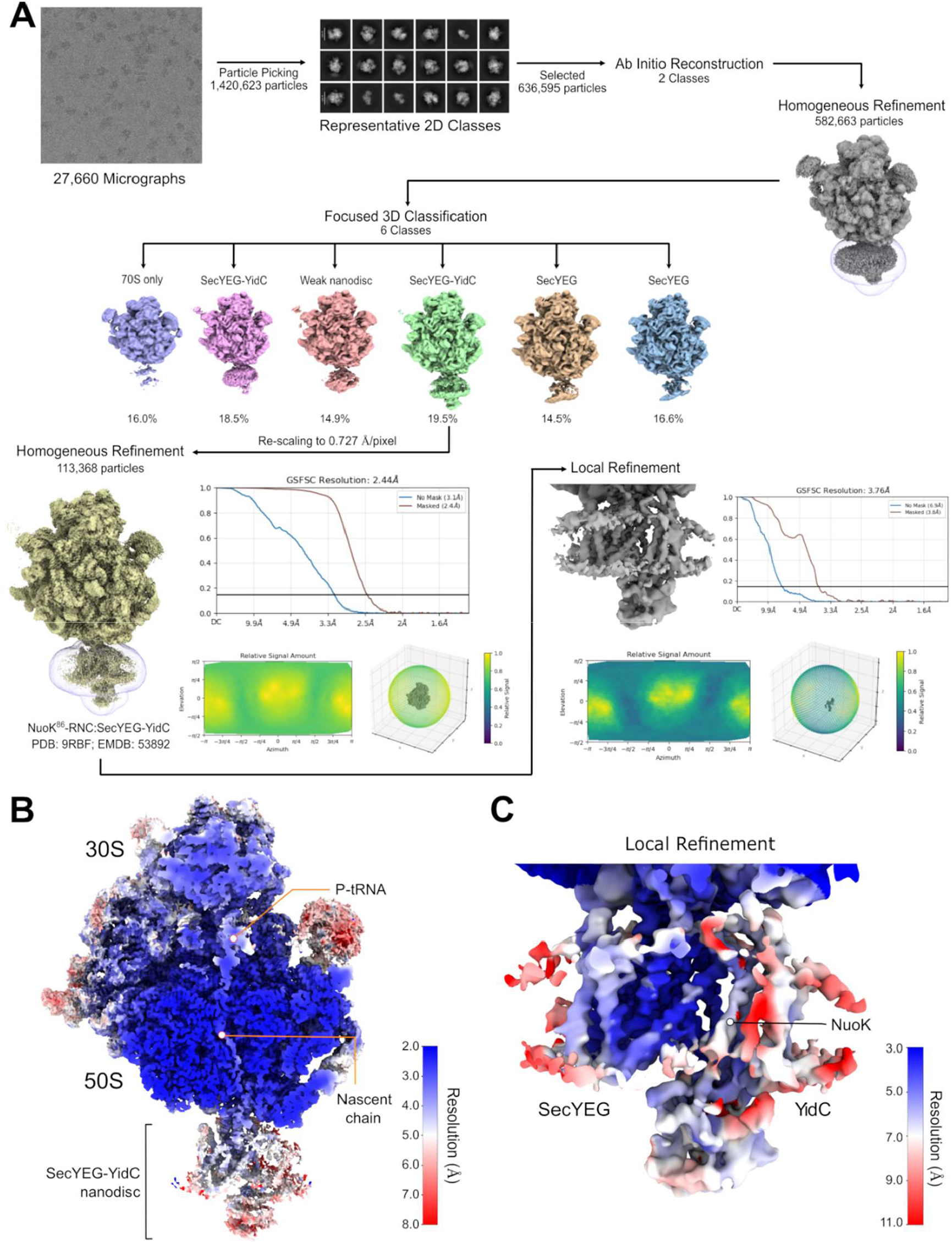
Cryo-EM data analysis, classification and resolution of the crosslinked NuoK^86^-RNC:SecYEG-YidC complex. **(A)** Classification and refinement scheme employed for single-particle analysis. The structurally distinct classes, distribution of views and Fourier shell correlation plots are shown. **(B)** Globally refined cryo-EM map for the main SecYEG-YidC class, with the color-code according to the local resolution. Gold-standard Fourier shell correlation (GS-FSC) curve as well as angular distribution plot are shown. **(C)** Locally refined cryo-EM map of the nanodisc-embedded insertase with the color-code according to the local resolution.

**Figure EV3.**
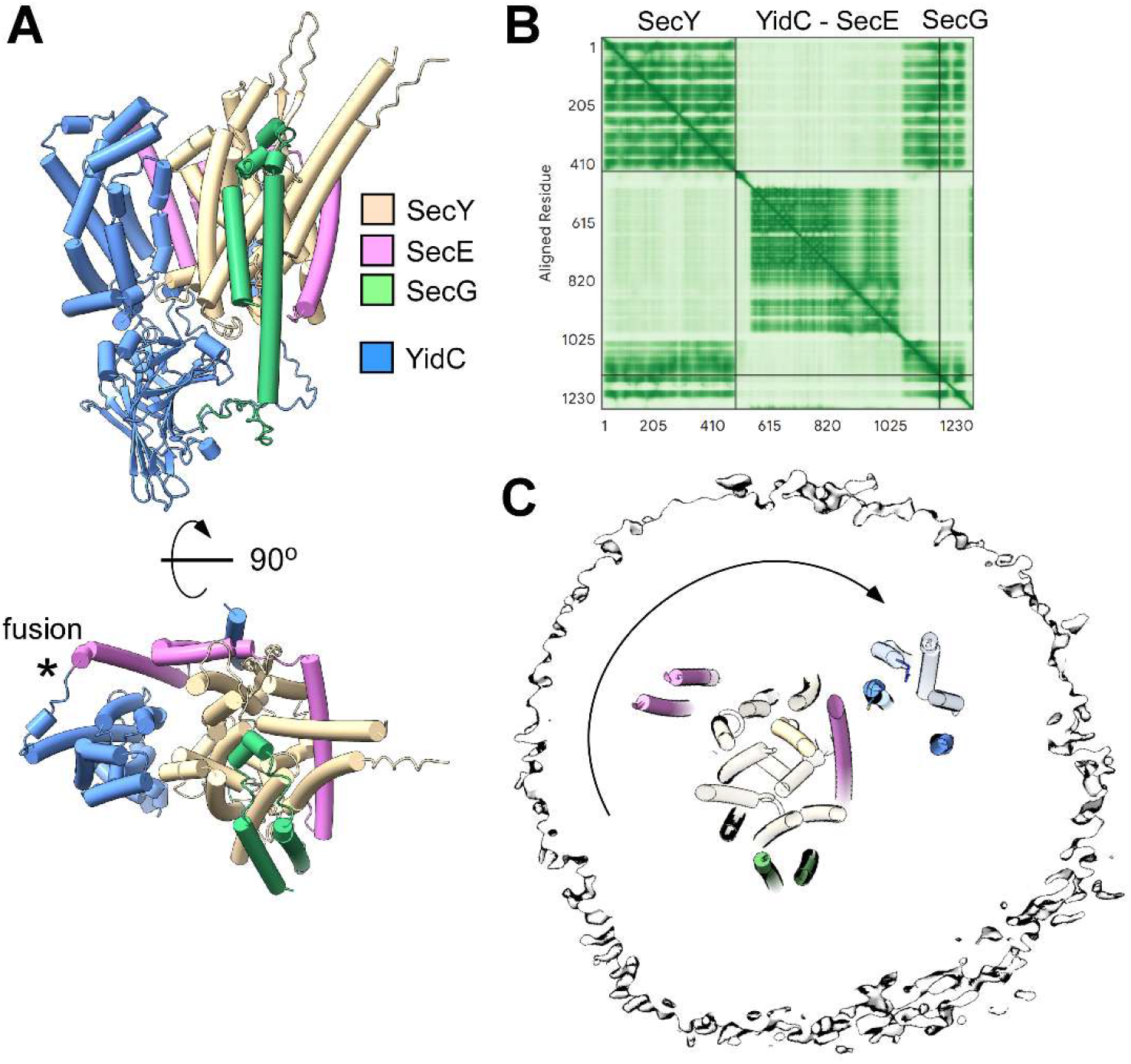
Repositioning of YidC within the nanodisc upon the assembly with SecYEG. **(A)** AlphaFold3-based model of SecYEG-YidC with the genetically fused YidC-SecE, as employed in reconstitution. YidC is positioned at the lateral gate. **(B)** PAE plot for the model shown in **(A)** suggests low confidence for SecY-YidC interactions. **(C)** Cross-section of MSP2N2 nanodisc shows the available lipid-filled space around SecYEG-YidC, allowing for YidC diffusion.

**Figure EV4.**
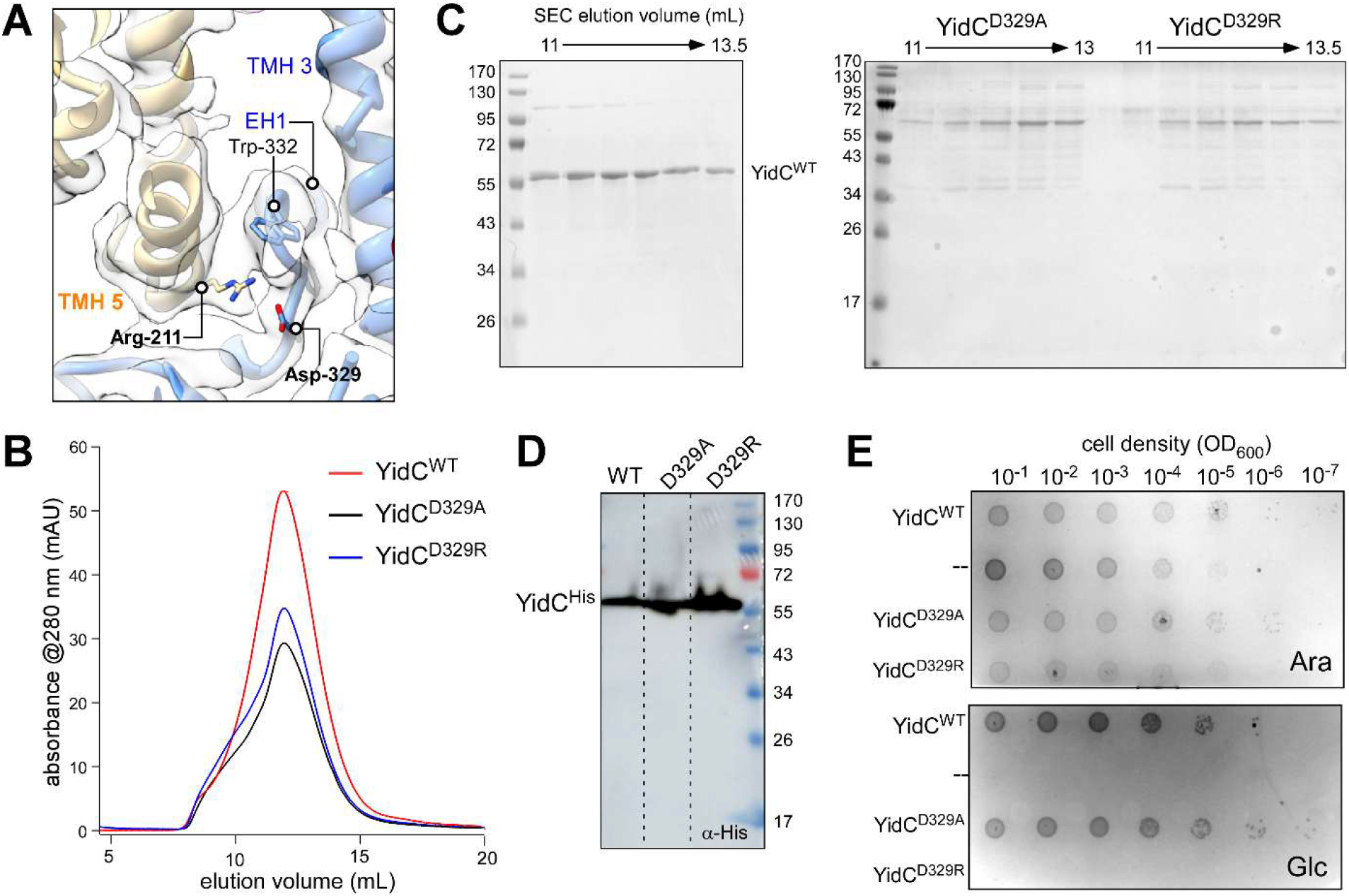
Charge inversion at YidC position 329 is deleterious for the cells. **(A)** View focusing on the periplasmic SecY-YidC interface displaying the molecular model of the SecY-YidC assembly fitted into the cryo-EM map (transparent). Proximate residues SecY Arg-211 and YidC Asp-329 are indicated in bold. **(B)** Size exclusion chromatography profiles of the DDM-solubilized and purified YidC variants. **(C)** SDS-PAGE of the size exclusion chromatography fractions shown in (**B**). **(D)** Immunodetection of the poly-histidine-tagged YidC variants expressed in *E. coli* FTL10 under permissive conditions (0.2% arabinose, 37 °C). **(E)** Complementation assay to test the effect of mutations at position 329 of YidC under permissive (0.2% arabinose, “Ara”) and the complementation (0.2 % glucose, “Glc”) conditions. The assay was performed at 37 °C.

**Figure EV5.**
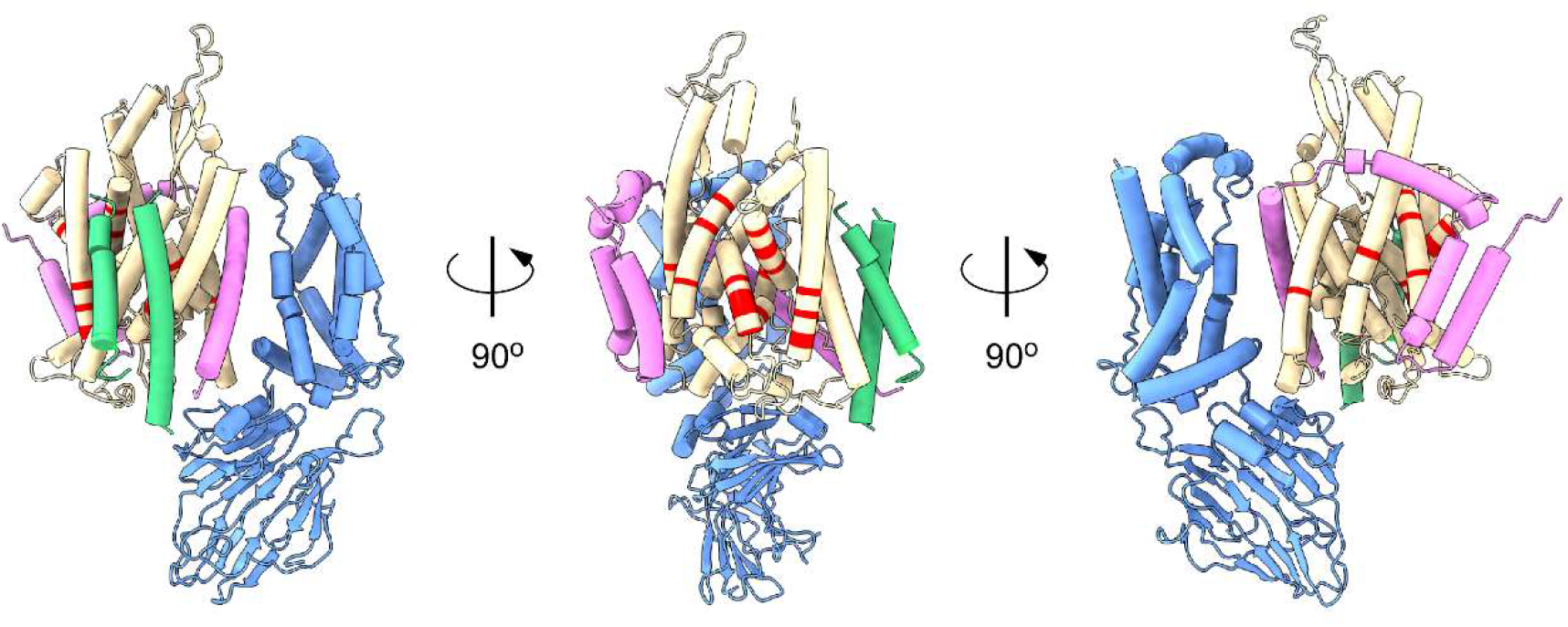
Overview of the site-specific crosslinking positions to probe SecYEG-YidC interactions. Positions employed to introduce photo-crosslinkers within SecYEG in earlier studies are shown in red (Jauss *et al*., 2019; Petriman *et al*., 2018; Sachelaru *et al*., 2013).

## References

Adams PD, Afonine PV, Bunkoczi G, Chen VB, Davis IW, Echols N, Headd JJ, Hung LW, Kapral GJ, Grosse-Kunstleve RW et al (2010) PHENIX: a comprehensive Python-based system for macromolecular structure solution. Acta Crystallogr D Biol Crystallogr 66: 213–221

Anghel SA, McGilvray PT, Hegde RS, Keenan RJ (2017) Identification of Oxa1 Homologs Operating in the Eukaryotic Endoplasmic Reticulum. Cell Rep 21: 3708–3716

Aviram N, Ast T, Costa EA, Arakel EC, Chuartzman SG, Jan CH, Hassdenteufel S, Dudek J, Jung M, Schorr S et al (2016) The SND proteins constitute an alternative targeting route to the endoplasmic reticulum. Nature 540: 134–138

Bischoff L, Wickles S, Berninghausen O, van der Sluis EO, Beckmann R (2014) Visualization of a polytopic membrane protein during SecY-mediated membrane insertion. Nat Commun 5: 4103

Botte M, Zaccai NR, Nijeholt JL, Martin R, Knoops K, Papai G, Zou J, Deniaud A, Karuppasamy M, Jiang Q et al (2016) A central cavity within the holo-translocon suggests a mechanism for membrane protein insertion. Sci Rep 6: 38399

Chen Y, Capponi S, Zhu L, Gellenbeck P, Freites JA, White SH, Dalbey RE (2017) YidC Insertase of Escherichia coli: Water Accessibility and Membrane Shaping. Structure 25: 1403–1414 e1403

Chiba K, Mori H, Ito K (2002) Roles of the C-terminal end of SecY in protein translocation and viability of Escherichia coli. J Bacteriol 184: 2243–2250

Cymer F, Hedman R, Ismail N, von Heijne G (2015a) Exploration of the arrest peptide sequence space reveals arrest-enhanced variants. J Biol Chem 290: 10208–10215

Cymer F, von Heijne G, White SH (2015b) Mechanisms of integral membrane protein insertion and folding. J Mol Biol 427: 999–1022

du Plessis DJ, Berrelkamp G, Nouwen N, Driessen AJ (2009) The lateral gate of SecYEG opens during protein translocation. J Biol Chem 284: 15805–15814

Emsley P, Cowtan K (2004) Coot: model-building tools for molecular graphics. Acta Crystallogr D Biol Crystallogr 60: 2126–2132

Gemmer M, Chaillet ML, van Loenhout J, Cuevas Arenas R, Vismpas D, Grollers-Mulderij M, Koh FA, Albanese P, Scheltema RA, Howes SC et al (2023) Visualization of translation and protein biogenesis at the ER membrane. Nature 614: 160–167

Gersteuer F, Morici M, Gabrielli S, Fujiwara K, Safdari HA, Paternoga H, Bock LV, Chiba S, Wilson DN (2024) The SecM arrest peptide traps a pre-peptide bond formation state of the ribosome. Nat Commun 15: 2431

Goddard TD, Huang CC, Meng EC, Pettersen EF, Couch GS, Morris JH, Ferrin TE (2018) UCSF ChimeraX: Meeting modern challenges in visualization and analysis. Protein Sci 27: 14–25

Haque ME, Spremulli LL, Fecko CJ (2010) Identification of protein-protein and protein-ribosome interacting regions of the C-terminal tail of human mitochondrial inner membrane protein Oxa1L. J Biol Chem 285: 34991–34998

Hatzixanthis K, Palmer T, Sargent F (2003) A subset of bacterial inner membrane proteins integrated by the twin-arginine translocase. Mol Microbiol 49: 1377–1390

Itskanov S, Park E (2023) Mechanism of Protein Translocation by the Sec61 Translocon Complex. Cold Spring Harb Perspect Biol 15: a041250

Jauss B, Petriman NA, Drepper F, Franz L, Sachelaru I, Welte T, Steinberg R, Warscheid B, Koch HG (2019) Noncompetitive binding of PpiD and YidC to the SecYEG translocon expands the global view on the SecYEG interactome in Escherichia coli. J Biol Chem 294: 19167–19183

Jomaa A, Boehringer D, Leibundgut M, Ban N (2016) Structures of the E. coli translating ribosome with SRP and its receptor and with the translocon. Nat Commun 7: 10471

Kalinin IA, Peled-Zehavi H, Barshap ABD, Tamari SA, Weiss Y, Nevo R, Fluman N (2024) Features of membrane protein sequence direct post-translational insertion. Nat Commun 15: 10198

Kanonenberg K, Royes J, Kedrov A, Poschmann G, Angius F, Solgadi A, Spitz O, Kleinschrodt D, Stuhler K, Miroux B et al (2019) Shaping the lipid composition of bacterial membranes for membrane protein production. Microb Cell Fact 18: 131

Kater L, Frieg B, Berninghausen O, Gohlke H, Beckmann R, Kedrov A (2019) Partially inserted nascent chain unzips the lateral gate of the Sec translocon. EMBO Rep 20: e48191

Kedrov A, Sustarsic M, de Keyzer J, Caumanns JJ, Wu ZC, Driessen AJ (2013) Elucidating the native architecture of the YidC: ribosome complex. J Mol Biol 425: 4112–4124

Kedrov A, Wickles S, Crevenna AH, van der Sluis EO, Buschauer R, Berninghausen O, Lamb DC, Beckmann R (2016) Structural Dynamics of the YidC:Ribosome Complex during Membrane Protein Biogenesis. Cell Rep 17: 2943–2954

Kempf N, Remes C, Ledesch R, Zuchner T, Hofig H, Ritter I, Katranidis A, Fitter J (2017) A Novel Method to Evaluate Ribosomal Performance in Cell-Free Protein Synthesis Systems. Sci Rep 7: 46753

Koch S, Driessen AJM, Kedrov A (2018) Biophysical Analysis of Sec-Mediated Protein Translocation in Nanodiscs. Adv Biomembr Lipid Self-Assem 28: 41–85

Kohler R, Boehringer D, Greber B, Bingel-Erlenmeyer R, Collinson I, Schaffitzel C, Ban N (2009) YidC and Oxa1 form dimeric insertion pores on the translating ribosome. Mol Cell 34: 344–353

Kumazaki K, Chiba S, Takemoto M, Furukawa A, Nishiyama K, Sugano Y, Mori T, Dohmae N, Hirata K, Nakada-Nakura Y et al (2014a) Structural basis of Sec-independent membrane protein insertion by YidC. Nature 509: 516–520

Kumazaki K, Kishimoto T, Furukawa A, Mori H, Tanaka Y, Dohmae N, Ishitani R, Tsukazaki T, Nureki O (2014b) Crystal structure of Escherichia coli YidC, a membrane protein chaperone and insertase. Sci Rep 4: 7299

Li Z, Boyd D, Reindl M, Goldberg MB (2014) Identification of YidC residues that define interactions with the Sec Apparatus. J Bacteriol 196: 367–377

Lu Y, Turnbull IR, Bragin A, Carveth K, Verkman AS, Skach WR (2000) Reorientation of aquaporin-1 topology during maturation in the endoplasmic reticulum. Mol Biol Cell 11: 2973–2985

McDowell MA, Heimes M, Sinning I (2021) Structural and molecular mechanisms for membrane protein biogenesis by the Oxa1 superfamily. Nat Struct Mol Biol 28: 234–239

Miyazaki R, Ai M, Tanaka N, Suzuki T, Dhomae N, Tsukazaki T, Akiyama Y, Mori H (2022) Inner membrane YfgM-PpiD heterodimer acts as a functional unit that associates with the SecY/E/G translocon and promotes protein translocation. J Biol Chem 298: 102572

Ou X, Ma C, Sun D, Xu J, Wang Y, Wu X, Wang D, Yang S, Gao N, Song C et al (2025) SecY translocon chaperones protein folding during membrane protein insertion. Cell 188: 1912–1924.e13

Page KR, Nguyen VN, Pleiner T, Tomaleri GP, Wang ML, Guna A, Hazu M, Wang TY, Chou TF, Voorhees RM (2024) Role of a holo-insertase complex in the biogenesis of biophysically diverse ER membrane proteins. Mol Cell 84: 3302–3319.e3311

Petriman NA, Jauss B, Hufnagel A, Franz L, Sachelaru I, Drepper F, Warscheid B, Koch HG (2018) The interaction network of the YidC insertase with the SecYEG translocon, SRP and the SRP receptor FtsY. Sci Rep 8: 578

Price CE, Driessen AJ (2008) YidC is involved in the biogenesis of anaerobic respiratory complexes in the inner membrane of Escherichia coli. J Biol Chem 283: 26921–26927

Price CE, Driessen AJM (2010) Conserved negative charges in the transmembrane segments of subunit K of the NADH:ubiquinone oxidoreductase determine its dependence on YidC for membrane insertion. J Biol Chem 285: 3575–3581

Punjani A, Rubinstein JL, Fleet DJ, Brubaker MA (2017) cryoSPARC: algorithms for rapid unsupervised cryo-EM structure determination. Nat Methods 14: 290–296

Ravaud S, Stjepanovic G, Wild K, Sinning I (2008) The crystal structure of the periplasmic domain of the Escherichia coli membrane protein insertase YidC contains a substrate binding cleft. J Biol Chem 283: 9350–9358

Ritchie TK, Grinkova YV, Bayburt TH, Denisov IG, Zolnerciks JK, Atkins WM, Sligar SG (2009) Chapter 11 - Reconstitution of membrane proteins in phospholipid bilayer nanodiscs. Methods Enzymol 464: 211–231

Sachelaru I, Petriman NA, Kudva R, Kuhn P, Welte T, Knapp B, Drepper F, Warscheid B, Koch HG (2013) YidC occupies the lateral gate of the SecYEG translocon and is sequentially displaced by a nascent membrane protein. J Biol Chem 288: 16295–16307

Seppala S, Slusky JS, Lloris-Garcera P, Rapp M, von Heijne G (2010) Control of membrane protein topology by a single C-terminal residue. Science 328: 1698–1700

Seurig M, Ek M, von Heijne G, Fluman N (2019) Dynamic membrane topology in an unassembled membrane protein. Nat Chem Biol 15: 945–948

Smalinskaite L, Hegde RS (2023) The Biogenesis of Multipass Membrane Proteins. Cold Spring Harb Perspect Biol 15: a041251

Smalinskaite L, Kim MK, Lewis AJO, Keenan RJ, Hegde RS (2022) Mechanism of an intramembrane chaperone for multipass membrane proteins. Nature 611: 161–166

Sohlenkamp C, Geiger O (2016) Bacterial membrane lipids: diversity in structures and pathways. FEMS Microbiol Rev 40: 133–159

Soman R, Yuan J, Kuhn A, Dalbey RE (2014) Polarity and charge of the periplasmic loop determine the YidC and sec translocase requirement for the M13 procoat lep protein. J Biol Chem 289: 1023–1032

Steinberg R, Knupffer L, Origi A, Asti R, Koch HG (2018) Co-translational protein targeting in bacteria. FEMS Microbiol Lett 365: fny095

Sundaram A, Li Q, Wan Y, Tang J, Wu H, Smalinskaite L, Hegde RS, Ji Z, Keenan RJ (2025) Global analysis of translocon remodeling during protein synthesis at the ER. Nat Struct Mol Biol 32: 2517–2525

Sundaram A, Yamsek M, Zhong F, Hooda Y, Hegde RS, Keenan RJ (2022) Substrate-driven assembly of a translocon for multipass membrane proteins. Nature 611: 167–172

Tsukazaki T (2018) Structure-based working model of SecDF, a proton-driven bacterial protein translocation factor. FEMS Microbiol Lett 365: fny112

Tsukazaki T (2019) Structural Basis of the Sec Translocon and YidC Revealed Through X-ray Crystallography. Protein J 38: 249–261

Urbanus ML, Froderberg L, Drew D, Bjork P, de Gier JW, Brunner J, Oudega B, Luirink J (2002) Targeting, insertion, and localization of Escherichia coli YidC. J Biol Chem 277: 12718–12723

von Heijne G (2007) The membrane protein universe: what’s out there and why bother? J Intern Med 261: 543–557

Voorhees RM, Hegde RS (2016) Structure of the Sec61 channel opened by a signal sequence. Science 351: 88–91

Watson ZL, Ward FR, Meheust R, Ad O, Schepartz A, Banfield JF, Cate JH (2020) Structure of the bacterial ribosome at 2 A resolution. Elife 9: e60482

White SH, Wimley WC (1999) Membrane protein folding and stability: physical principles. Annu Rev Biophys Biomol Struct 28: 319–365

Wickles S, Singharoy A, Andreani J, Seemayer S, Bischoff L, Berninghausen O, Soeding J, Schulten K, van der Sluis EO, Beckmann R (2014) A structural model of the active ribosome-bound membrane protein insertase YidC. Elife 3: e03035

Wirth JO, Scheiderer L, Engelhardt T, Engelhardt J, Matthias J, Hell SW (2023) MINFLUX dissects the unimpeded walking of kinesin-1. Science 379: 1004–1010

Woodall NB, Hadley S, Yin Y, Bowie JU (2017) Complete topology inversion can be part of normal membrane protein biogenesis. Protein Sci 26: 824–833

Wu H, Smalinskaite L, Hegde RS (2024) EMC rectifies the topology of multipass membrane proteins. Nat Struct Mol Biol 31: 32–41

Wu ZC, de Keyzer J, Kedrov A, Driessen AJ (2012) Competitive binding of the SecA ATPase and ribosomes to the SecYEG translocon. J Biol Chem 287: 7885–7895

Yang TJ, Mukherjee S, Langer JD, Hummer G, McDowell MA (2025) SND3 is the membrane insertase within a distinct SEC61 translocon complex. Nat Commun 16: 9566

Yuan J, Phillips GJ, Dalbey RE (2007) Isolation of cold-sensitive yidC mutants provides insights into the substrate profile of the YidC insertase and the importance of transmembrane 3 in YidC function. J Bacteriol 189: 8961–8972

Zheng SQ, Palovcak E, Armache JP, Verba KA, Cheng Y, Agard DA (2017) MotionCor2: anisotropic correction of beam-induced motion for improved cryo-electron microscopy. Nat Methods 14: 331–332

Zhu L, Kaback HR, Dalbey RE (2013) YidC protein, a molecular chaperone for LacY protein folding via the SecYEG protein machinery. J Biol Chem 288: 28180–28194

Zubay G (1973) In vitro synthesis of protein in microbial systems. Annu Rev Genet 7: 267–287

