## Supplemental Figures for "Substrate-induced assembly and functional mechanism of the bacterial membrane protein insertase SecYEG-YidC"

### Appendix Figures

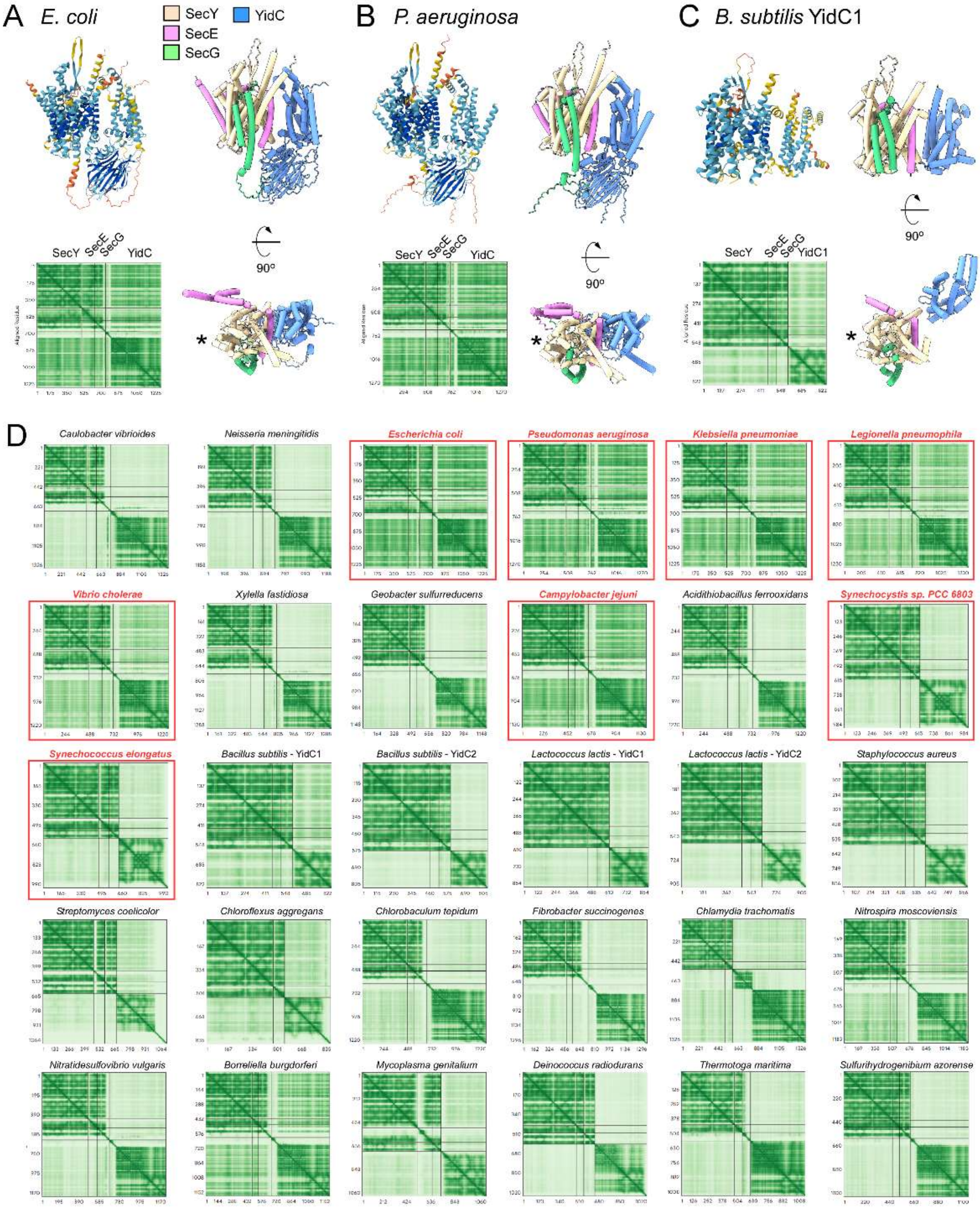

**Appendix Figure S1. Diverse architectures of modelled SecYEG-YidC complexes.** AlphaFold3-based prediction for SecYEG-YidC assembly for *Escherichia coli* **(A)**, *Pseudomonas aeruginosa* **(B)**, and *Bacillus subtilis* with the primary YidC homolog (YidC1/SpoIIIJ) **(C).** The uniform color coding of the proteins is indicated in **(A).** The position of the SecY lateral gate for each model is indicated with the asterisk. **(D)** Predicted aligned error (PAE) plots for SecYEG-YidC models from indicated bacterial species (see also Suppl. Table 1). Representatives from the major bacterial phyla were chosen based on a recent phylogeny analysis (Hug *et al.* 2016 *Nature Microbiol*). The models matching the cryo-EM structure of *E. coli* SecYEG-YidC complex are indicated in red. The generated AlphaFold models are provided in the Dataset EV2.

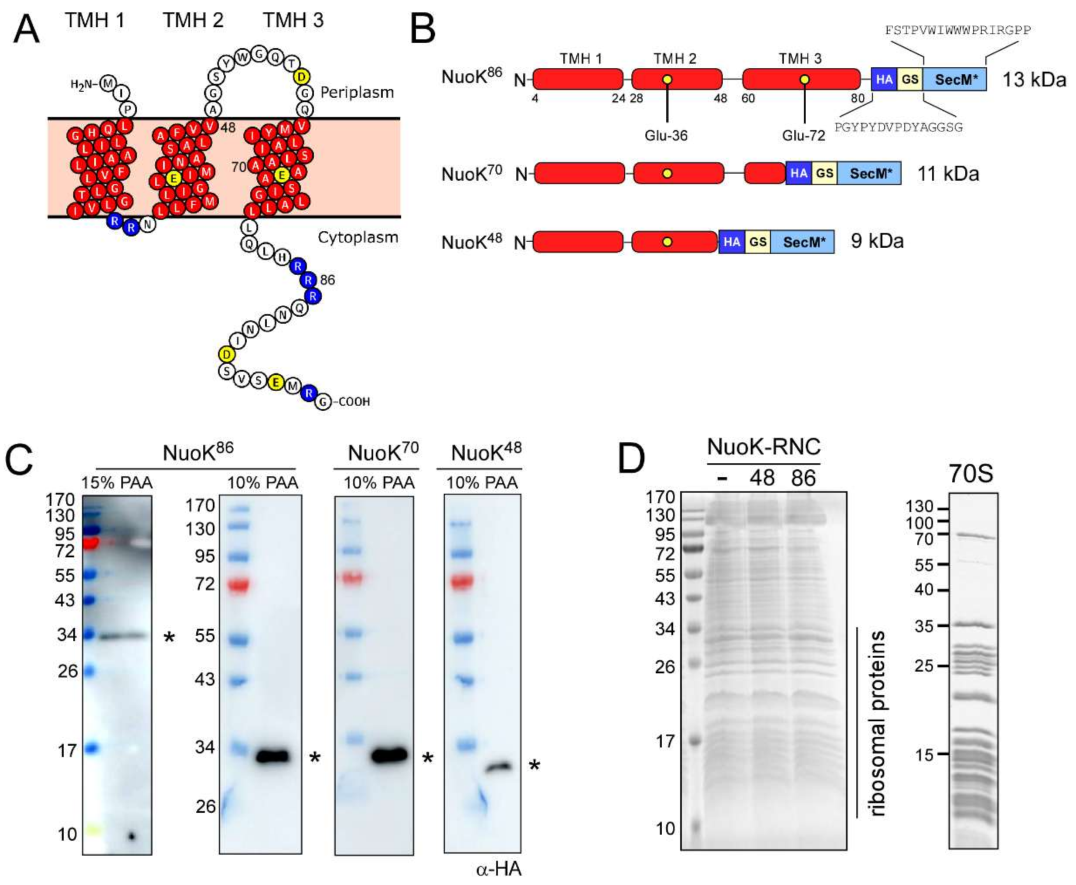

**Appendix Figure S2. Design of the NuoK nascent chain constructs.**

(**A**) Primary structure of *E. coli* NuoK. Residues within TMHs are shown in red, charged residues are shown in blue (arginines) and yellow (aspartates, glutamates). Terminal residues of the tested nascent chains (Val-48, Ala-70 and Arg-86) are indicated by their numbers. The image is prepared via Protter service (wlab.ethz.ch/protter) and modified in Inkscape (Inkscape Project).

(**B**) Design of the nascent chains mimicking NuoK$^{48}$ and NuoK$^{86}$ insertion intermediates. NuoK TMHs are shown in red with the flanking residue positions indicated. Positions of the glutamate residues within TMHs 2 and 3 are shown as yellow circles. The sequence of the C-terminal linker consisting of the HA-tag (HA) and a glycine-serine linker (GS) is shown for NuoK$^{86}$. SecM* represents a SecM-based ribosome stalling sequence.

(**C**) Western blots against the HA-tag on the isolated NuoK-RNC:SecYEG-YidC complexes. The NuoK nascent chains coupled to tRNA (additional molecular mass ~25 kDa) are indicated with asterisks. For NuoK$^{86}$-RNC blots were performed using SDS-PAGE with either 15 or 10 % polyacrylamide (PAA), as indicated.

(**D**) SDS-PAGE of *E. coli* membrane isolates after stalling NuoK translation *in vivo* showing equal amounts of the total protein content, incl. the abundant ribosomal proteins. A characteristic SDS-PAGE of purified 70S ribosomes is provided as a reference.

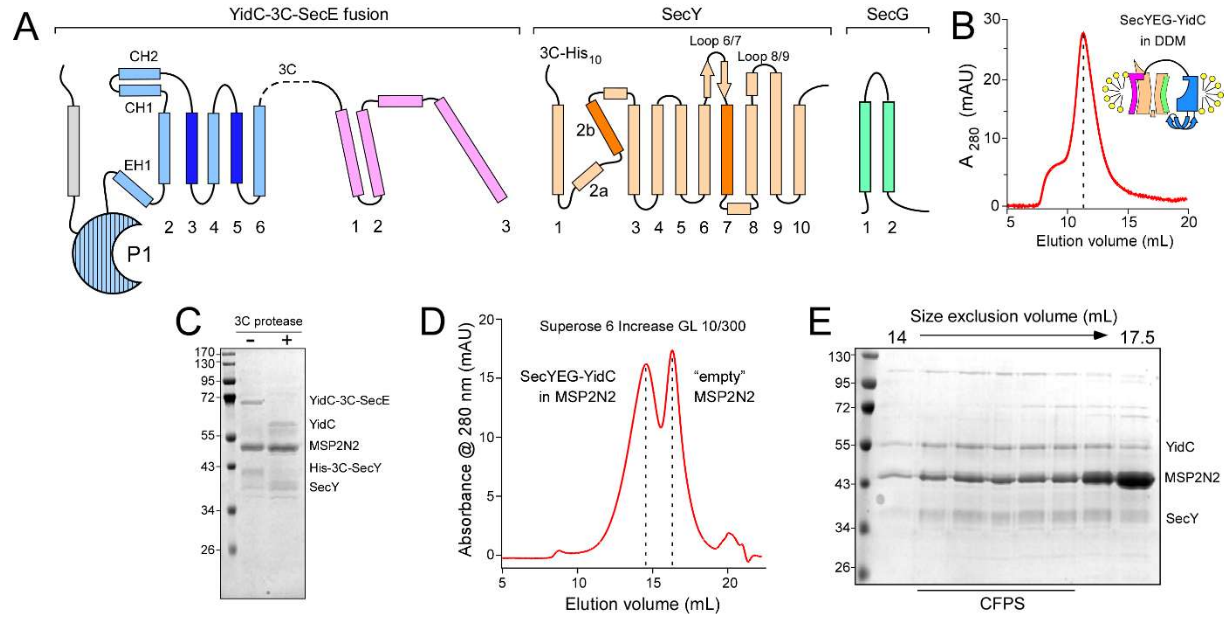

**Appendix Figure S3. Preparation of the nanodisc-reconstituted SecYEG-YidC.**

(**A**) The architecture of the SecYEG-YidC fusion complex. YidC TMHs 3 and 5 forming the insertion site are shown in dark blue, SecY TMHs 2b and 7 forming the lateral gate are shown in dark orange. The non-essential YidC TMH 1 (not resolved in the cryo-EM map) is shown in grey.

(**B**) Size exclusion chromatography profile of the detergent-isolated SecYEG-YidC complex.

(**C**) SDS-PAGE of the nanodisc-reconstituted SecYEG-YidC complex before and after HRV-3C protease treatment.

(**D**) Size exclusion chromatography profile of the nanodisc reconstitution reaction. Peaks corresponding to MSP2N2-nanodiscs containing SecYEG-YidC and those containing only lipids ("empty") are indicated with the dashed lines.

(**E**) SDS-PAGE of the size exclusion chromatography fractions shown in (**D**).

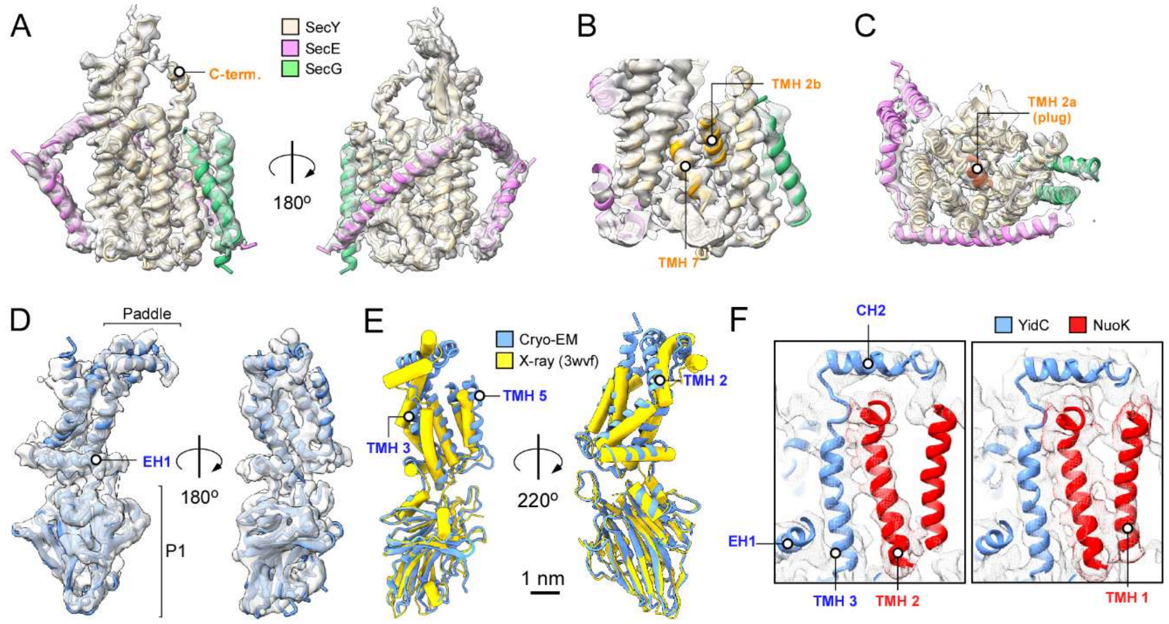

**Appendix Figure S4. Architecture of the SecYEG-YidC insertase and the substrate NuoK.**

(**A**) Side views on the isolated cryo-EM density (transparent) of SecYEG with fitted model. The C-terminal extension of SecY is indicated (C-term).

(**B**) View on the lateral gate of SecYEG formed by SecY TMHs 2b and 7.

(**C**) View on SecYEG from the periplasmic side showing the central sealing position of the "plug" domain, TMH 2a.

(**D**) Side views on the isolated cryo-EM density of YidC with the fitted model. Selected structural elements are indicated.

(**E**) An overlay of the cryo-EM-based model of YidC within the active insertase complex (blue ribbons) with the X-ray structure of the idle YidC (PDB ID 3WVF; yellow cylinders). TMHs with changed positions are indicated.

(**F**) Putative position of the membrane-inserted NuoK TMH 1. A rod-shaped density is revealed at a lower contour level (right).

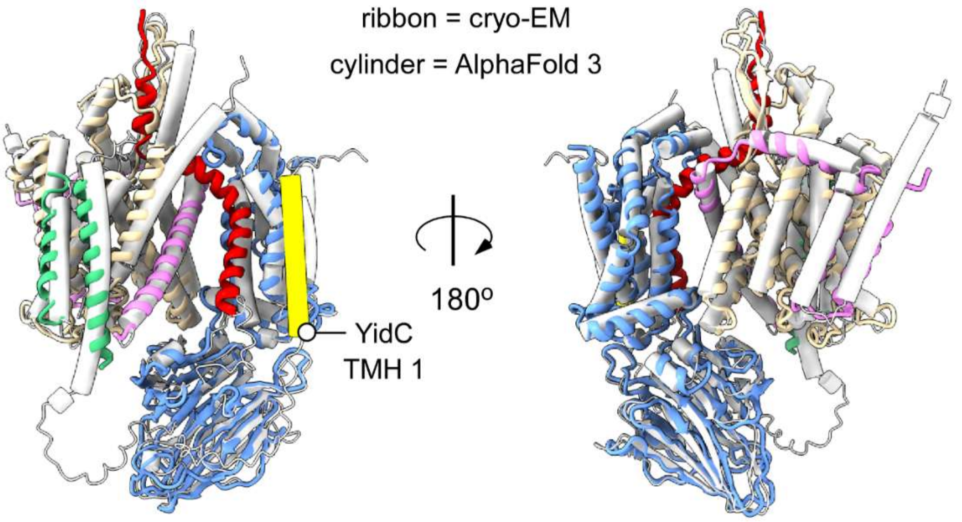

**Appendix Figure S5. Computational model vs resolved structure of SecYEG-YidC complex.**

Comparison of the cryo-EM-derived model for the SecYEG-YidC complex in the presence of NuoK86-RNC (ribbon representation) and the AlphaFold 3-based model (cylinder representation).

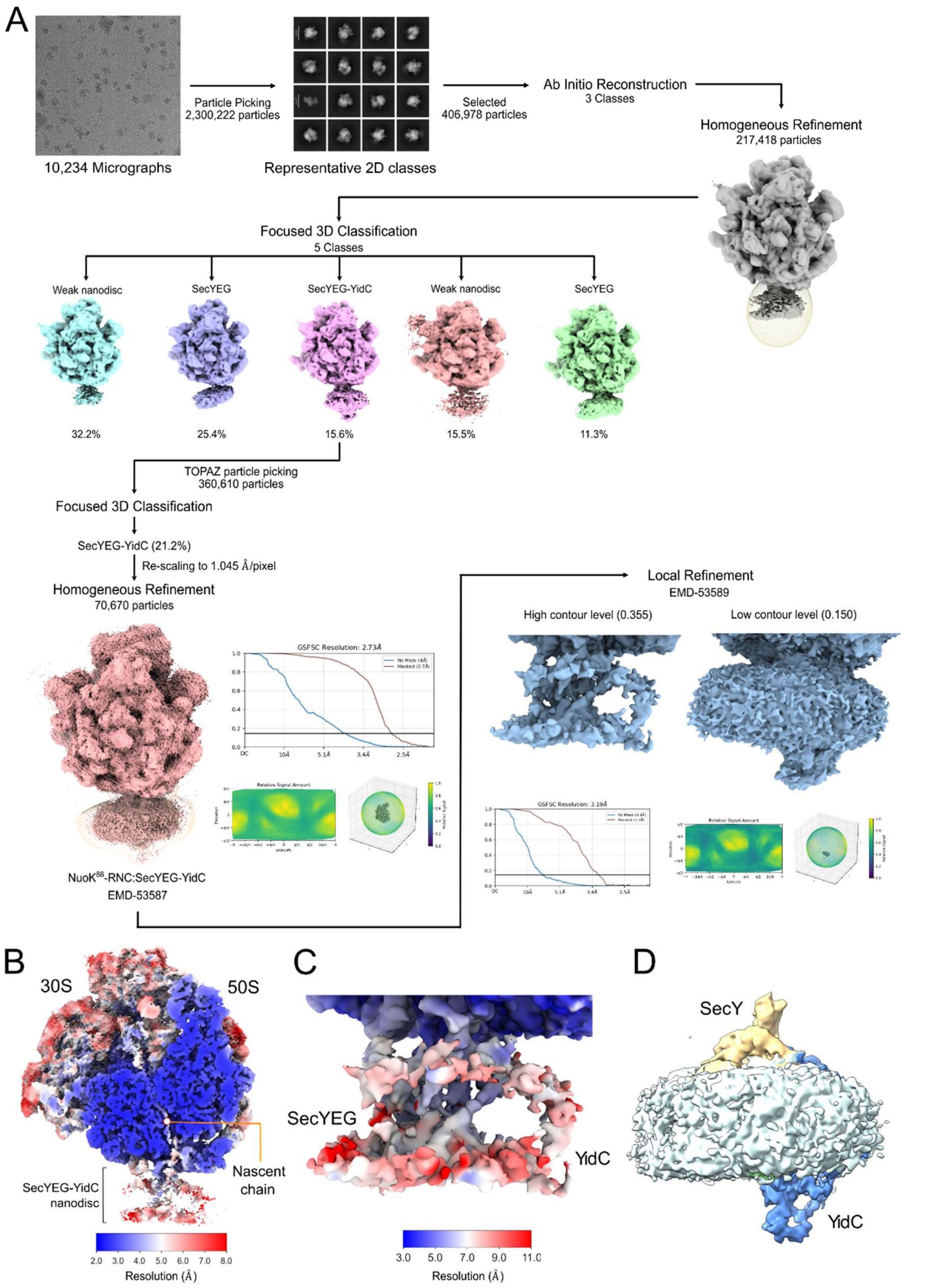

**Appendix Figure S6. Cryo-EM data analysis, classification and resolution of the non-crosslinked NuoK$^{86}$-RNC:SecYEG-YidC complex.**

**(A)** From a total of 10,234 micrographs, 2,300,222 particles were picked and used for 2D classification using cryoSPARC, which yielded a total of 408,798 ribosomal particles (70S) with an extra density at the tunnel exit. After *ab-initio* reconstruction and homogenous refinement, 217,418 particles were selected for focused 3D classification with a mask covering the tunnel exit density (shown in transparent yellow). This yielded five classes, of which one class showed density for SecYEG and YidC including the P1 domain. This class was used to train a TOPAZ picking model. 360,610 particles were classified and refined again, yielding a final map at 2.73 Å average resolution from 70,670 particles. Local refinement on the SecYEG-YidC region resulted in map of 3.19 Å resolution that showed density for YidC at low contour levels. Gold-standard Fourier Shell Correlation (GS-FSC) curves as well as angular distribution plots are shown for globally and locally refined maps.

**(B, C)** Globally and locally refined cryo-EM maps are color-coded according to local resolution.

**(D)** Cryo-EM reconstruction of the non-crosslinked NuoK$^{86}$-RNC:SecYEG-YidC complex. The focused refined map was colored based on the molecular model of the crosslinked SecYEG-YidC complex (Figure 2A). The extra density for YidC was present at the “back-of-Sec” position suggesting that the architecture of SecYEG-YidC complex was not affected by the stabilizing crosslinking procedure.

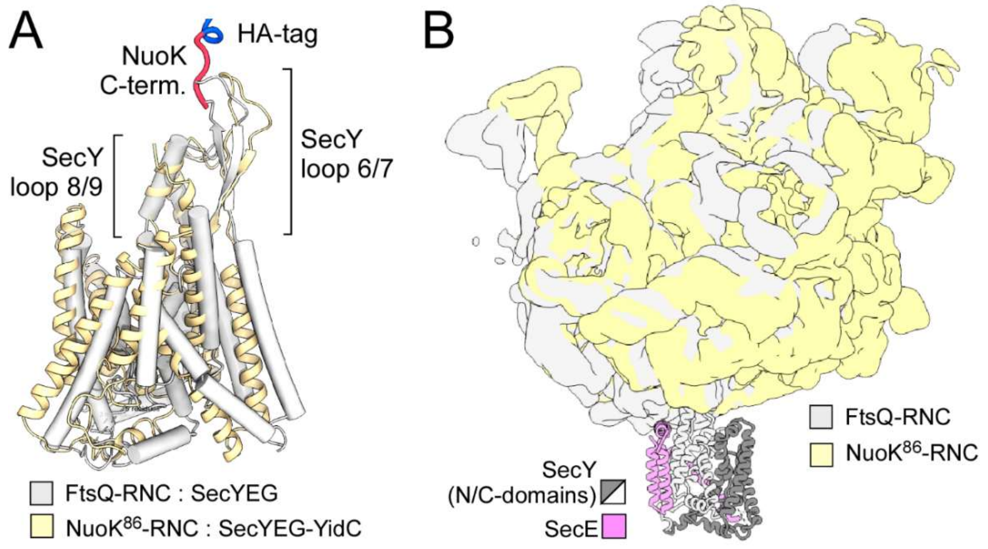

**Appendix Figure S7. Docking geometry of SecYEG on RNC depends on the nascent chain.**

**(A)** Orientation of SecY bound to NuoK$^{86}$-RNC within the SecYEG-YidC complex (ribbon representation) and that bound to FtsQ-RNC (PDB ID 6R7L, cylinder representation) (aligned on the 50S ribosomal subunit).

**(B)** Orientations of FtsQ-RNC and NuoK$^{86}$-RNC (aligned on the SecYEG complex). The 50S ribosomal subunit in the NuoK$^{86}$-RNC is tilted towards the N-terminal half of SecY (grey).

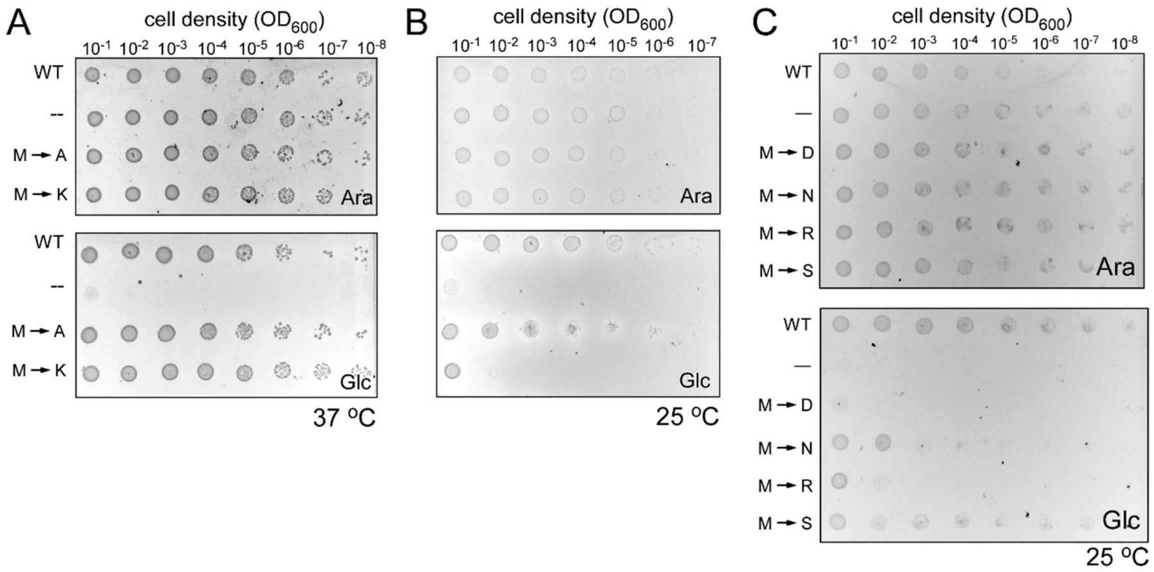

**Appendix Figure S8. Mutations in the YidC paddle domain affect *E. coli* viability at low temperature.** Complementation assay to test the effect of mutations within the paddle domain of YidC (substitution of Met-408/409 by indicated amino acids) under permissive (0.2% arabinose, "Ara") and the complementation (0.2 % glucose, "Glc") conditions. The assay was performed at 37 °C **(A)** and at 25 °C **(B, C).**

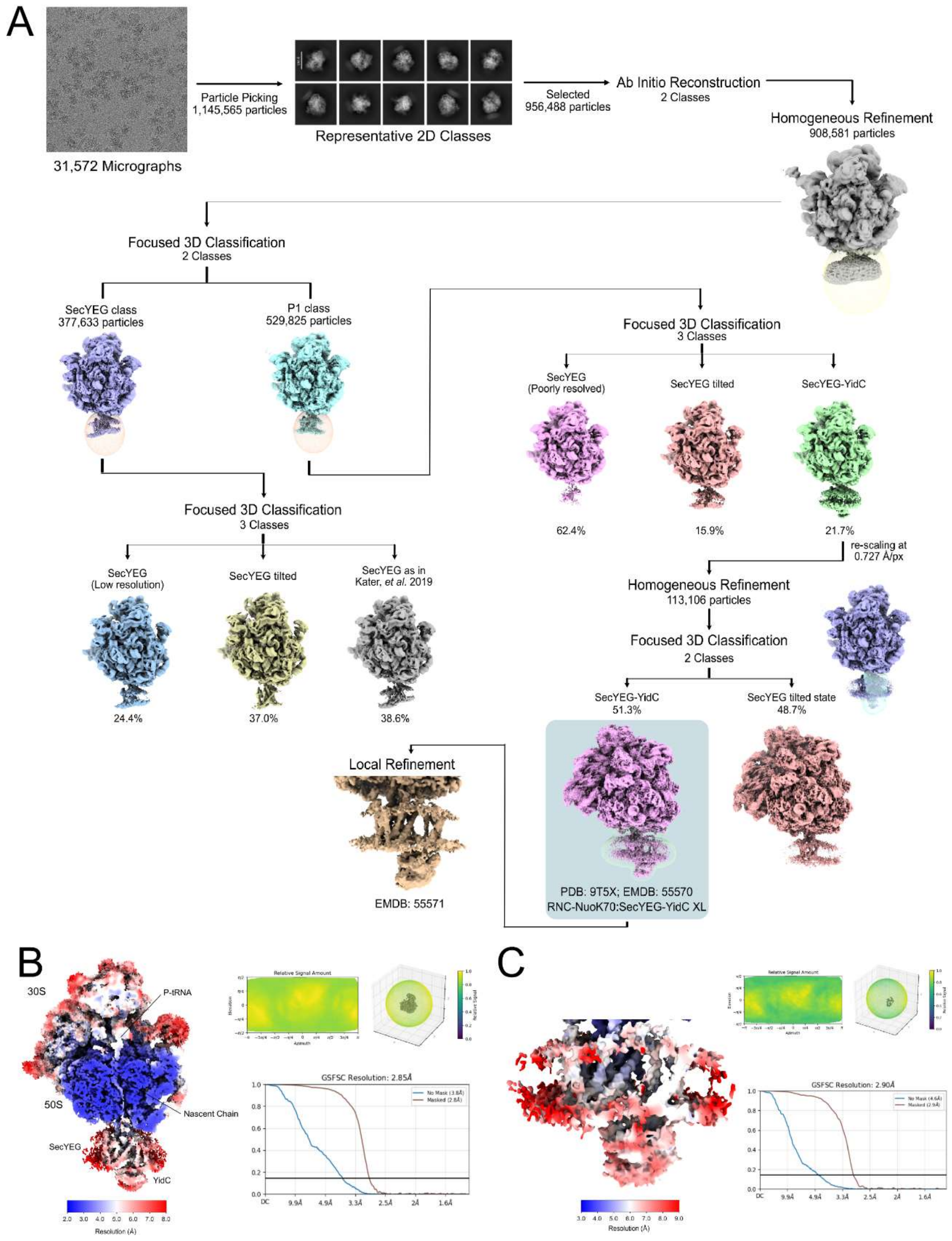

**Appendix Figure S9. Cryo-EM data analysis, classification and resolution of the NuoK$^{70}$-RNC:SecYEG-YidC complex.**

**(A)** From a total of 31,572 micrographs, 956,488 particles were selected after 2D classification using cryoSPARC, which yielded a total of 908,581 ribosomal particles (70S) with an extra density at the tunnel exit. Several rounds of 3D focused classification using masks around the tunnel exit/nanodisc region yielded one class with SecYEG and YidC present and two well-resolved classes with only SecYEG present, one in the conformation as observed in Kater et al., one in the tilted conformation as observed in presence of YidC (see also Appendix Fig. 7). The SecY-YidC-containing map was refined to a final resolution of 2.85 Å average resolution. Local refinement on the SecYEG-YidC region resulted in map of 2.90 Å resolution.

(**B, C**) Globally and locally refined cryo-EM maps shown color coded according to local resolution. Gold-standard Fourier Shell Correlation (GS-FSC) curves as well as angular distribution plots are shown for globally and locally refined maps.

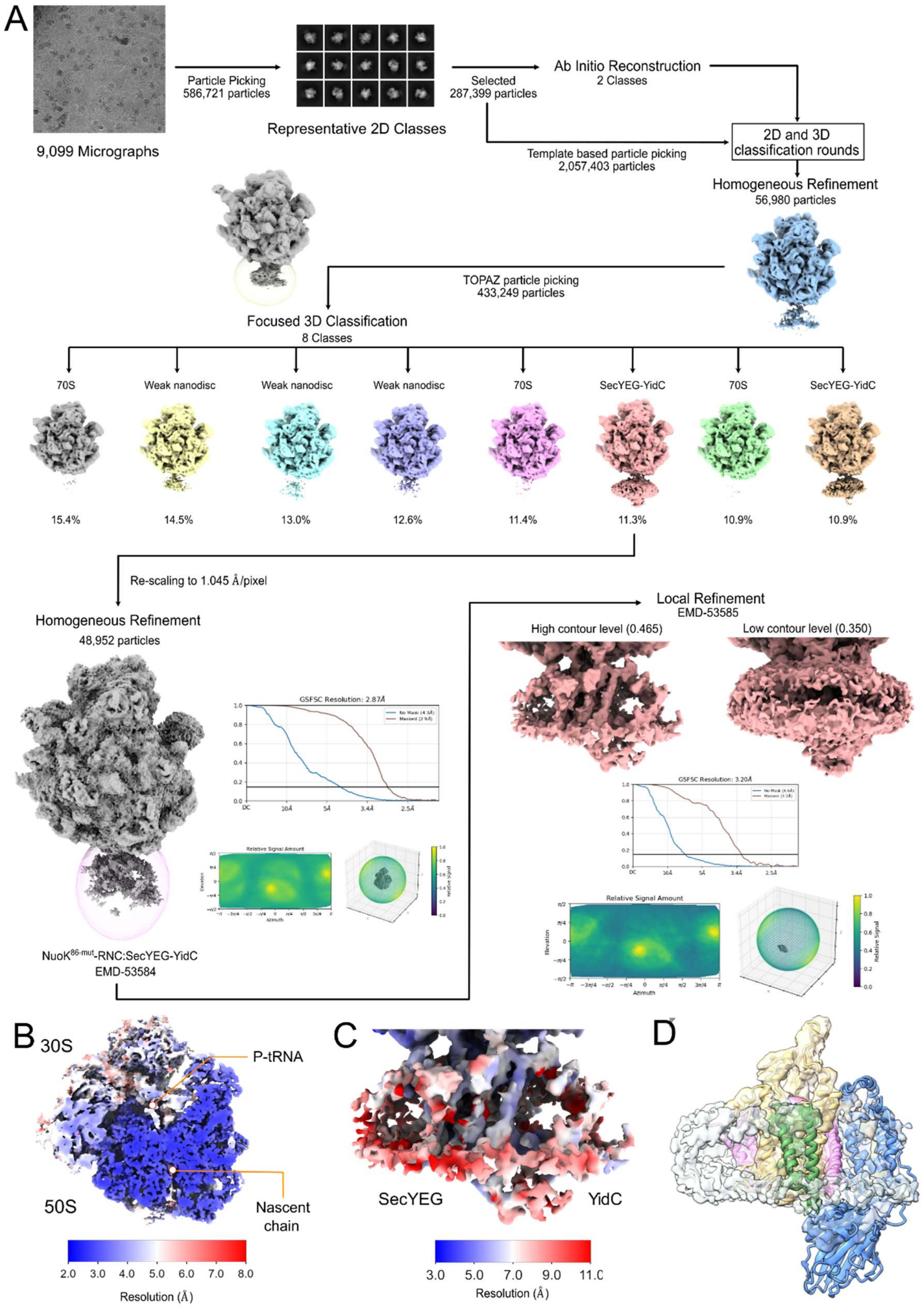

**Appendix Figure S10. Cryo-EM data analysis, classification and resolution of the NuoK$^{86mut}$-RNC:SecYEG-YidC complex.**

(**A**) From a total of 9,099 micrographs, 586,721 particles were picked and used for 2D classification using cryoSPARC, which yielded a total of 287,399 (70S) ribosomal particles with an extra density at the tunnel exit. A similar procedure to optimize particle picking and 3D classification was performed as for the non-crosslinked NuoK$^{86}$-RNC:SecYEG-YidC complex (Appendix Fig. S6) yielding a class with 48,952 particles with well-defined density for SecYEG and YidC including the P1 domain below the tunnel exit. This class was refined to a final average resolution of 2.87 Å. Local refinement on the SecYEG/YidC region resulted in map of 3.20 Å resolution that showed density for YidC at low contour levels. Gold-standard Fourier Shell Correlation (GS-FSC) curves as well as angular distribution plots are shown for globally and locally refined maps.

(**B, C**) Globally and locally refined cryo-EM maps shown color coded according to local resolution.

(**D**) Cryo-EM reconstruction of the NuoK86mut-RNC:SecYEG-YidC complex. The focused refined map was colored based on the molecular model of the crosslinked SecYEG-YidC complex (Figure 2A). Note that also here extra density for YidC was present at the “back-of Sec” position suggesting that the architecture of SecYEG-YidC complex was not affected by the E36K, E72K double mutation in the NuoK nascent chain.

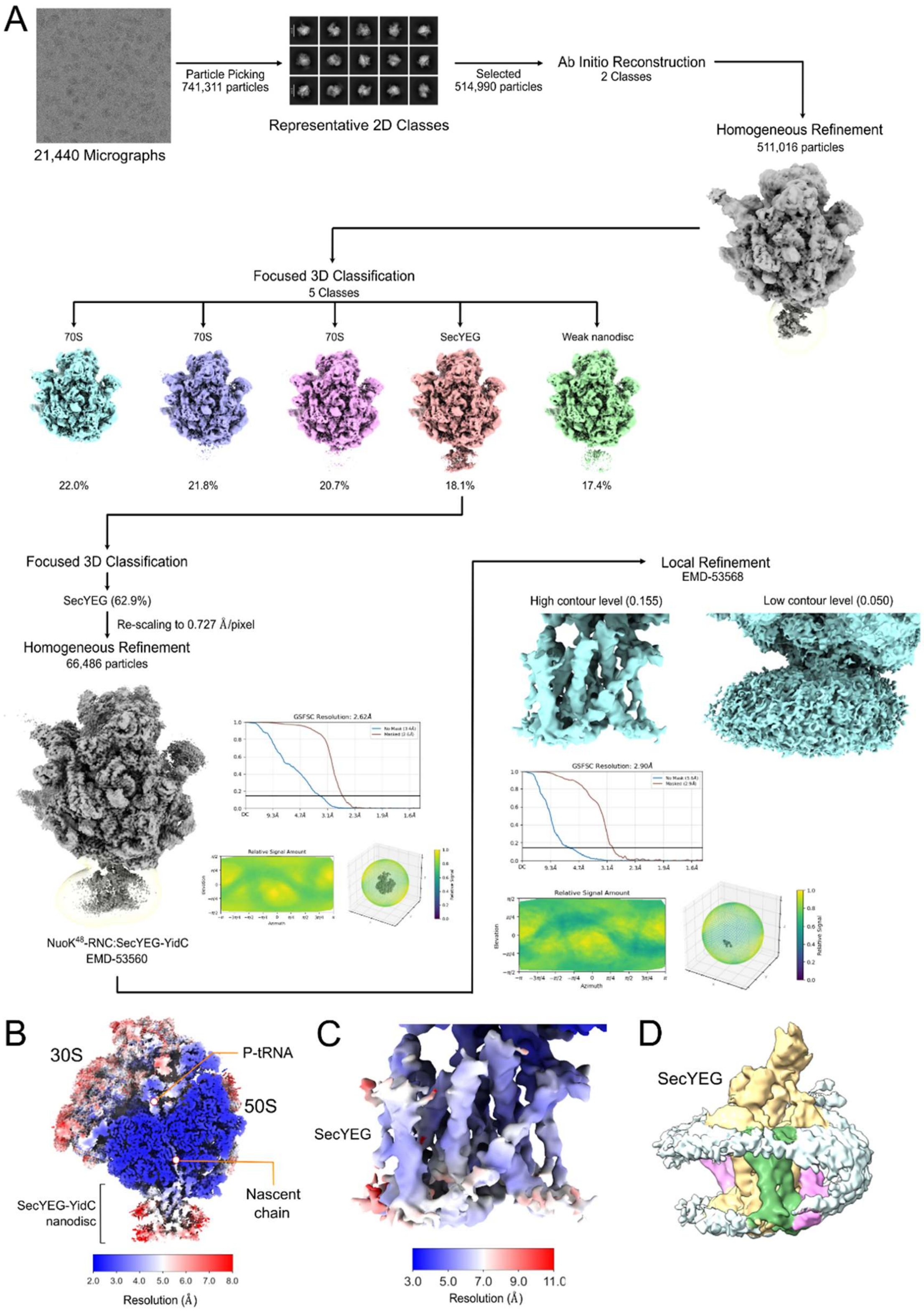

**Appendix Figure S11. Cryo-EM data analysis, classification and resolution of the NuoK$^{48}$-RNC:SecYEG complex.**

(**A**) From a total of 21,440 micrographs, 741,311 particles were picked and used for 2D classification using cryoSPARC, which yielded a total of 514,990 ribosomal particles (70S) with an extra density at the tunnel exit. This dataset was essentially process as outlined for the NuoK$^{86}$-RNC:SecYEG-

YidC dataset (Figure EV2). Here, focused 3D classification only yielded classes with no ligand or only SecYEG bound to the tunnel exit. The best-resolved class with 66,486 particles was refined to a final average resolution of 2.62 Å. Local refinement on the Nanodisc/SecYEG region resulted in map of 2.90 Å resolution that shown all α-helices of SecYEG clearly resolved. Gold-standard Fourier Shell Correlation (GS-FSC) curves as well as angular distribution plots are shown for globally and locally refined maps.

(**B, C**) Globally and locally refined cryo-EM maps shown color coded according to local resolution.

(**D**) Cryo-EM reconstruction of the $NuoK^{48}$-RNC:SecYEG complex. The focused refined map was colored based on the molecular model of the SecYEG complex (Figure 2A). No extra density for YidC was present at the “back-of-Sec” position suggesting that YidC is not stably recruited.

**Appendix Table S1. AlphaFold 3 predictions of SecYEG-YidC complexes of different bacterial clades**

| No. | Bacterial clade | Species | Position relative to SecYEG | ipTM | pTM | Comments |
|---|---|---|---|---|---|---|
| 1 | Alphaproteobacterium | *Caulobacter vibrioides* | SecG | 0,44 | 0,48 | |
| 2 | Betaproteobacterium | *Neisseria meningitidis* | SecG | 0,45 | 0,49 | |
| 3 | Gammaproteobacterium | *Escherichia coli* | Back of Sec | 0,69 | 0,71 | |
| 4 | Gammaproteobacterium | *Pseudomonas aeruginosa* | Back of Sec | 0,69 | 0,71 | |
| 5 | Gammaproteobacterium | *Klebsiella pneumoniae* | Back of Sec | 0,63 | 0,66 | |
| 6 | Gammaproteobacterium | *Legionella pneumophila Lens* | Back of Sec | 0,6 | 0,63 | |
| 7 | Gammaproteobacterium | *Vibrio cholerae* | Back of Sec | 0,55 | 0,58 | |
| 8 | Gammaproteobacterium | *Xylella fastidiosa* | SecG | 0,45 | 0,5 | |
| 9 | Deltaproteobacterium | *Geobacter sulfurreducens* | SecG | 0,45 | 0,49 | |
| 10 | Epsilonproteobacterium/ Campylobacterota | *Campylobacter jejuni* | Back of Sec | 0,63 | 0,66 | |
| 11 | Acidithiobacillus class | *Acidithiobacillus ferrooxidans* | SecG | 0,45 | 0,5 | |
| 12 | Cyanobacteria | *Synechocystis sp. PCC 6803* | Back of Sec | 0,5 | 0,53 | |
| 13 | Cyanobacteria | *Synechococcus elongatus PCC 7942* | Back of Sec | 0,51 | 0,55 | |
| 14 | Firmicutes | *Bacillus subtilis - YidC1* | Back of Sec | 0,59 | 0,63 | YidC is rotated ~180 grad |
| 15 | Firmicutes | *Bacillus subtilis - YidC2* | SecG | 0,53 | 0,59 | |
| 16 | Firmicutes | *Lactococcus lactis - YidC1* | SecG | 0,53 | 0,61 | |
| 17 | Firmicutes | *Lactococcus lactis - YidC2* | SecG | 0,51 | 0,58 | |
| 18 | Firmicutes | *Staphylococcus aureus* | SecY TMH 6/TMH7 | 0,52 | 0,56 | |
| 19 | Actinobacteria | *Streptomyces coelicolor* | Lateral gate | 0,52 | 0,55 | |
| 20 | Chloroflexi | *Chloroflexus aggregans* | Lateral gate | 0,5 | 0,56 | |
| 21 | Chlorobi | *Chlorobaculum tepidum* | Lateral gate | 0,44 | 0,48 | Model #1 (correct topology) |
| 22 | Fibrobacteres | *Fibrobacter succinogenes* | Lateral gate | 0,43 | 0,47 | |
| 23 | PVC — Chlamydiae | *Chlamydia trachomatis serovar D* | Back of Sec | 0,42 | 0,47 | YidC is rotated ~90 grad |
| 24 | Nitrospirae | *Nitrospira moscoviensis* | Inverted/non-physiological topology | 0,52 | 0,54 | |
| 25 | Thermodesulfobacteriota | *Nitratidesulfovibrio vulgaris* | SecG | 0,46 | 0,5 | |
| 26 | Spirochaetes | *Borreliella burgdorferi* | Back of Sec | 0,6 | 0,63 | YidC is rotated ~180 grad |
| 27 | Tenericutes / Mollicutes | *Mycoplasma genitalium* | Lateral gate | 0,5 | 0,54 | |
| 28 | Deinococcus–Thermus | *Deinococcus radiodurans* | SecG | 0,47 | 0,51 | |
| 29 | Thermotogae | *Thermotoga maritima* | Lateral gate | 0,4 | 0,46 | |
| 30 | Aquificota | *Sulfurihydrogenibium azorense* | SecG | 0,47 | 0,51 | Model #2 (correct topology) |

**Appendix Table S2. Cryo-EM data collection, refinement and validation statistics**

| | NuoK$^{86}$-RNC: SecYEG-YidC (crosslinked) (EMD-53892)[a] (EMD-53893)[b] (EMD-53894)[c] (PDB 9RBF) | NuoK$^{70}$-RNC: SecYEG-YidC (crosslinked) (EMD-55570)[a] (EMD-55571)[b] (EMD-55598)[c] (PDB-9T5X) | NuoK$^{86}$-RNC: SecYEG-YidC (EMD-53587)[a] (EMD-53589)[b] | NuoK$^{86mut}$-RNC: SecYEG-YidC (EMD-53584)[a] (EMD-53585)[b] | NuoK$^{48}$-RNC: SecYEG-YidC (EMD-53560)[a] (EMD-53568)[b] |
|---|---|---|---|---|---|
| **Data collection and processing** | | | | | |
| Magnification | 165,000 | 165,000 | 130,000 | 130,000 | 165,000 |
| Voltage (kV) | 300 | 300 | 300 | 300 | 300 |
| Electron exposure (e–/Å$^2$) | 60 | 60 | 40 | 40 | 60 |
| Defocus range (μm) | 0.5-3.5 | 0.5-3.5 | 0.5-3.5 | 0.5-3.5 | 0.5-3.5 |
| Pixel size (Å) | 0.727 | 0.727 | 1.060 | 1.049 | 0.727 |
| Symmetry imposed | C1 | C1 | C1 | C1 | C1 |
| Initial particle images (no.) | 1,420,623 | 1,145,565 | 2,300,222 | 2,057,403 | 741,311 |
| Final particle images (no.) | 113,368 | 58,063 | 70,670 | 48,952 | 66,486 |
| Map resolution (Å) | | | | | |
| FSC threshold (0.143) | 2.44[a], 3.76[b] | 2.85[a], 2.90[b] | 2.73[a], 3.19[b] | 2.87[a], 3.20[b] | 2.62[a], 2.90[b] |
| **Refinement** | | | | | |
| Initial model used (PDB code) | 7k00, 8qoa, AlphaFold | 9RBF | | | |
| Model resolution (Å) | 3.0 | 3.0 | | | |
| FSC threshold (0.5) | | | | | |
| Model composition | | | | | |
| Non-hydrogen atoms | 150,739 | 150,806 | | | |
| Protein residues | 6735 | 6723 | | | |
| Nucleotides | 4556 | 4556 | | | |
| Ligands | ZN, MG, SPM | ZN, MG, SPM | | | |
| *B* factors (mean) (Å$^2$) | | | | | |
| Protein | 87.07 | 187.83 | | | |
| Nucleotide | 72.51 | 152.70 | | | |
| Ligand | 49.64 | 105.46 | | | |
| R.m.s. deviations | | | | | |
| Bond lengths (Å) | 0.006 | 0.005 | | | |
| Bond angles (°) | 0.728 | 0.597 | | | |
| Validation | | | | | |
| MolProbity score | 1.33 | 1.09 | | | |
| Clashscore | 3.37 | 1.02 | | | |
| Poor rotamers (%) | 0.42 | 0.16 | | | |
| Ramachandran plot | | | | | |
| Favored (%) | 96.79 | 95.79 | | | |
| Allowed (%) | 3.15 | 4.18 | | | |
| Disallowed (%) | 0.06 | 0.03 | | | |

[a] Consensus refinement

[b] Local refinement

[c] Composite map
